# AP-1 Primes Transcriptional Noise-Led Cancer Treatment Adaptation Through KDM5 Chromatin Rewiring

**DOI:** 10.1101/2025.10.13.682004

**Authors:** Alejandro Allo Anido, Cecilia Roux, Ayeh Sadat Sadr, Siân Hamer, Emilia Chen, Abigail Shea, Christie English, Charlotte Butterworth, Harvey Che, Angella Bellini, Birgit Geoerger, Gudrun Schleiermacher, Louis Chesler, Michael David Morgan, Alejandra Bruna

## Abstract

How drug-tolerant persister (DTP) cells escape quiescence to drive tumour relapse remains a central unresolved question in cancer evolution. Here, we use neuroblastoma and hepatoblastoma, paediatric cancers with low genetic instability, as tractable model systems to study the principles and molecular mechanisms key in the early stages of adaptation to treatment. We identify transcriptional noise, the stochastic variability in gene expression, as a latent cellular property amplified post-treatment, during awakening from persistence, which drives adaptive regrowth following therapy withdrawal.

Through functional assays, lineage tracing and single-cell multiomics, we show that treatment enriches for progenitor-like tolerant states with limited detectable clonal selection. Upon drug withdrawal, awakening proceeds via a stochastic process driven by noise-enabled plasticity in cell identity programmes.

Mechanistically, AP-1-bound chromatin states emerge post-treatment, and AP-1 activity primes KDM5A recruitment to cell-state identity genes during awakening, driving H3K4me3 peak narrowing at their promoters and an increase in transcriptional noise that enables MES-to-ADRN plasticity. Pharmacological or genetic inhibition of AP-1 or KDM5 activity suppresses transcriptional noise and phenotypic plasticity, blocks DTP exit and prevents tumour recovery in both neuroblastoma and hepatoblastoma models.

Together, these results identify KDM5-mediated transcriptional noise, operating within AP-engaged chromatin landscapes, as a key driver of early adaptation to treatment. Targeting this axis offers a strategy to exploit DTP awakening as an evolutionary bottleneck to prevent treatment adaptation and cancer relapse.

**GRAPHICAL ABSTRACT:** 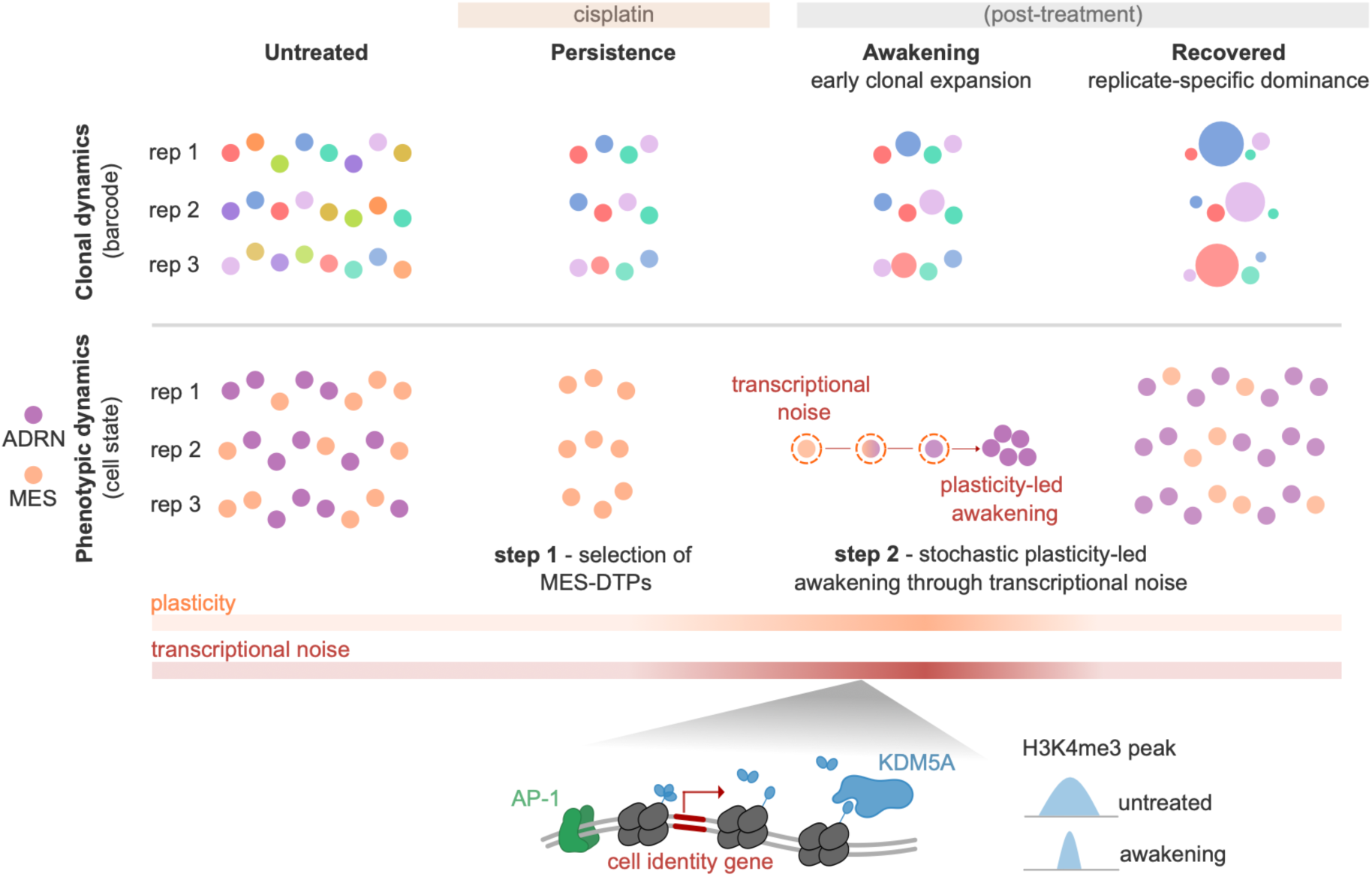

## INTRODUCTION

Cancer is the leading cause of death in children worldwide, accounting for more than one in five deaths among individuals under fourteen in the United Kingdom^1^. Unlike adult cancers, which often arise from an accumulation of somatic mutations, paediatric cancers typically emerge *in utero* due to disrupted differentiation and uncontrolled growth of transformed embryonic cells^2^. These malignancies are driven by relatively few genetic alterations^3,4^, yet they also frequently evade treatment due to their adaptive evolution capabilities. The embryonic cells from which paediatric cancers arise are inherently plastic during development, a property essential for their ability to differentiate into diverse cell types^5–9^. The developmental origin of childhood cancers, coupled with their low mutational burden, suggests that non-genetic mechanisms, rather than the extensive acquisition of novel mutations, play a key role in treatment resistance and relapse.

Phenotypic plasticity is the ability of cells to undergo dynamic and non-genetic cell state changes. Recently, it has been recognized as an emerging hallmark of cancer, where it amplifies heterogeneity and promotes treatment adaptation^10,11^. This phenomenon is tightly associated with chromatin rewiring^12,13^. One of the most prominent examples of phenotypic plasticity – both in development and cancer – is epithelial-to-mesenchymal transitions (EMT), in which cells switch from an epithelial transcriptional programme (apical-basal polarity, cell-cell contacts) to an invasive mesenchymal identity (front-rear polarity, migratory potential)^14^. Epigenetic changes also enable cancer progression, for instance, through the transient acquisition of stem cell potential in certain tumour subpopulations^15,16^. Additionally, epigenetically driven plasticity promotes treatment evasion in multiple cancers by enabling the entry into drug-tolerant persister (DTP) states: slow-cycling cells that are able to survive during treatment and have the potential to restart tumour proliferation upon treatment withdrawal^17^.

Cell fate decisions during development are often affected by a stochastic component of gene expression, known as transcriptional noise^18^. Transcriptional noise arises from intrinsic stochastic fluctuations in the process of gene expression within individual cells. It is primarily driven by the burst-like kinetics of transcription, in which promoters switch stochastically between active and inactive states, leading to sporadic mRNA production^18,19^. This bursting behaviour reflects fundamental molecular properties – such as promoter architecture, chromatin accessibility, transcription factor binding dynamics, and histone modification states – that influence the frequency and amplitude of transcriptional events^19^. While transcriptional noise describes the probabilistic behaviour of gene regulation within a single cell, it becomes experimentally observable as increased expression variability at the population level^20–22^. Such variability has been associated with phenotype diversification and cell-fate determination in multiple organisms, and it is considered a “bet-hedging” strategy to ensure survival in rapidly changing environments. Transcriptional noise facilitates HIV exit from latency^23^, promotes a competent phenotype in bacteria ^24^, contributes to organ development in mouse and the fruit fly^25,26^, and has more recently been linked to cancer progression and treatment resistance^27,28^. We therefore hypothesized that transcriptional noise may drive plasticity and adaptive evolution in paediatric cancers by increasing the likelihood of cells to spontaneously rewire their transcriptional programmes towards fitter phenotypes.

The advent of single-cell RNA sequencing has enabled unprecedented resolution in studying phenotypic dynamics in paediatric cancers. However, plasticity is inherently dynamic, and static single-cell snapshots lack the temporal resolution to disentangle true plastic transitions from fluctuations or selection of pre-existing states^29–31^. Here, we use a dynamic preclinical treatment framework in neuroblastoma and hepatoblastoma cell lines and patient-derived models to study at high temporal and molecular resolution the early evolutionary principles and molecular mechanisms underlying treatment adaptation.

By integrating lineage barcoding and single-cell multiomics, we show that treatment enriches for drug-tolerant persister states with cell cycle arrest and progenitor-like features. However, this phenotypic selection occurs with limited detectable clonal selection under the resolution of our barcode system.

Instead, upon drug withdrawal, the persister clones awaken stochastically through probabilistic cell state transitions into proliferative, more differentiated states that drive tumour regrowth. Post-treatment cell states display rewired chromatin with enriched accessibility in the vicinity of AP-1 motifs, a feature also seen in patient-derived post-treatment states that are linked to poor prognosis. AP-1 activity primes for KDM5A recruitment and local H3K4me3 peak narrowing at regulatory loci of cell identity genes, leading to an increase in transcriptional noise and phenotypic transitions from progenitor-like to differentiated-like cell states during awakening. Both AP-1 and KDM5A activities are essential for this plasticity-led post-treatment awakening and regrowth.

Collectively, our findings link chromatin remodelling to transcriptional noise and stochastic cell identity transitions, which drive post-treatment awakening and recovery. This positions transcriptional noise–driven plasticity as a key force in early cancer evolution and identifies KDM5A as an actionable regulator of adaptation and relapse in paediatric tumours.

## RESULTS

### Therapy selects phenotypes, but awakening is driven by phenotypic plasticity and stochastic clonal dynamics

Neuroblastoma, a tumour of the developing sympathetic nervous system, is the most common solid cancer in children after brain tumours^1^. Half of neuroblastoma patients present with “high-risk” disease and half of these will die due to treatment resistance or relapse^32,33^. As a paradigm of paediatric cancer, neuroblastoma exhibits limited genetic drivers, usually restricted to MYCN amplification and/or structural alterations in chromosome 17 and 1 (accounting to up to 65% high-risk patients). Other common drivers include TERT, ATRX and ALK ^34–36^. Genomic profiling of relapsed tumours has revealed further acquisition of alterations in these loci, but lack of shared novel drivers^37^, highlighting the contribution of non-genetic mechanisms to neuroblastoma evolution in patients. Despite this, neuroblastoma displays remarkable phenotypic heterogeneity. Two cell identities have been reported and well characterized in neuroblastoma pre-clinical models^38^. The noradrenergic identity (ADRN) – resembling differentiated neuroblasts – and the mesenchymal state (MES) – associated with treatment resistance and which some groups have linked to dedifferentiated Schwann cell precursors^39^. These ADRN and MES states are regulated by exclusive sets of transcription factors that form core regulatory circuitries (CRC)^40–42^. Recent studies support plasticity between ADRN and MES identities in pre-clinical models^29^, yet its impact on patients remains debatable. While some works question the presence of MES cells in patient tumours^39,43,44^, others suggest they only emerge in response to treatment^40,42^. Thus, studying the phenotypic dynamics and mechanisms driving phenotypic plasticity during treatment is essential to understand the processes fuelling resistance and relapse.

To unravel the contribution of plasticity to neuroblastoma evolution, we applied an expressed DNA barcoding approach within a preclinical treatment framework (Figure 1A). Individual SK-N-SH cells were marked with unique genomic barcodes detectable in both DNA and RNA sequencing platforms, enabling simultaneous tracking of clonal and phenotypic dynamics (Methods). In this system, one barcode is stably integrated into the genome and inherited across cell divisions, and therefore each barcode uniquely marks one lineage.

**Figure 1.**
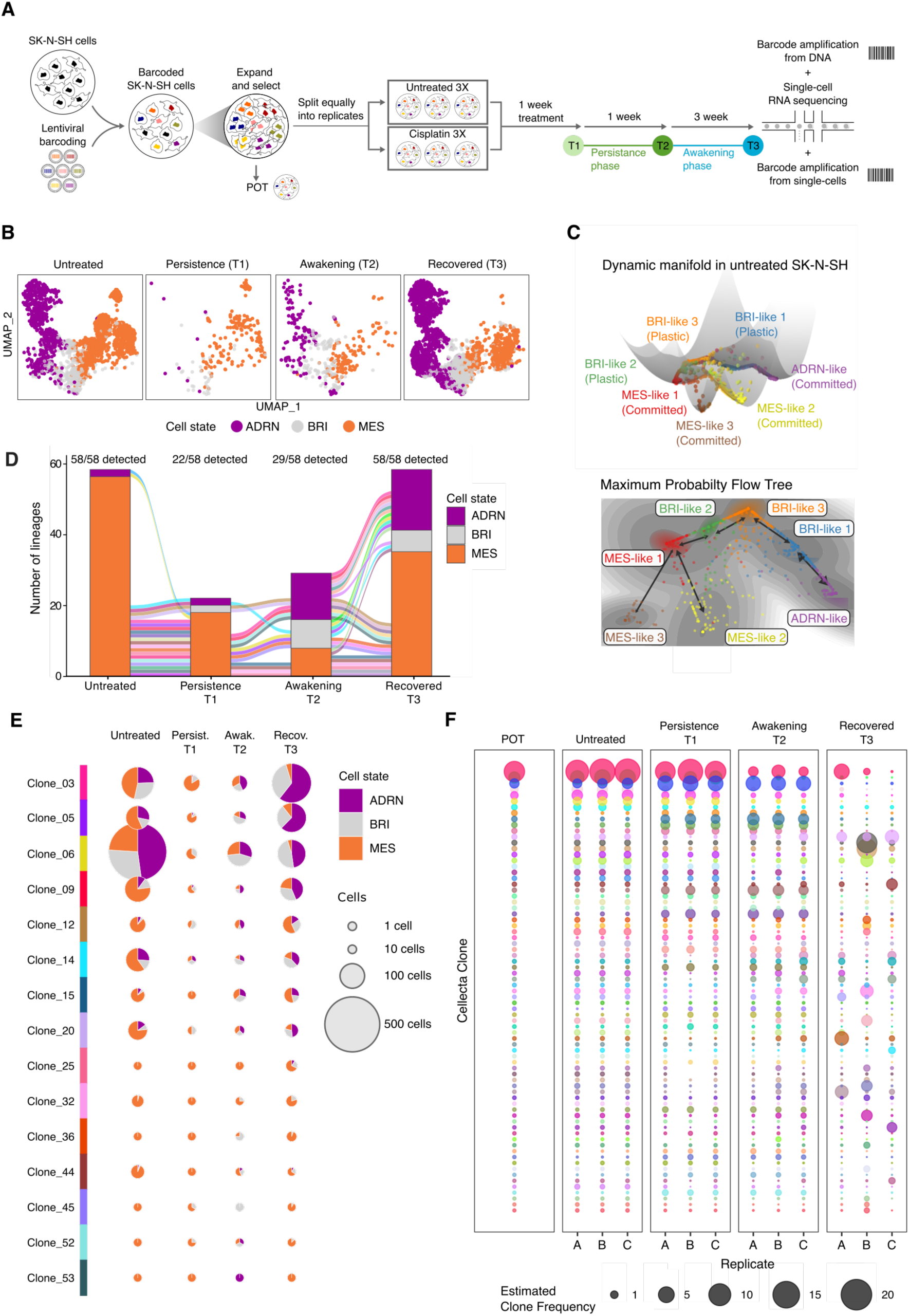
Lineage tracing reveals phenotypic selection followed by stochastic awakening of DTP cells in response to treatment. (A) SK-N-SH cells were tagged with Cellecta barcodes in a lentiviral system and expanded. The population (referred to as POT) was divided into triplicates and from each of these, the cells were distributed equally into six flasks. Among the six replicates, three were treated with IC50 doses of cisplatin, and the remaining three replicates served as untreated controls. After one week of treatment, 10% of the cells from each replicate were collected and cryopreserved (T1). The remaining cells in all replicates were re-seeded and left untreated. The collection process was repeated one week (T2) and four weeks (T4) later. (B) Umap of cell state occupancy (ADRN, adrenergic; BRI, bridge; MES, mesenchymal) across timepoints, derived from scRNA-seq data. (C) Dynamic manifold landscapes in untreated SK-N-SH cells. Well depth correlated with stability of the attractor state. Cells are coloured by attractor well and annotated as “plastic” or “committed” based on the entropy levels in (Figure S1B-C). Arrow size represents probability of transition between attractor wells. (D) Phenotypic dynamics of the 58 awakened lineages. Awakened lineages were defined as those detected in both untreated and recovered (T3) populations, and were assigned to their most abundant cell state. Coloured lines connect the same lineage in two timepoints. When a lineage disappears between timepoints, no cells containing that barcode were sampled in that specific timepoint. (E) Phenotypic composition of the 15 lineages shared across all treatment stages. (F) Frequency of the 100 most highly abundant lineages within POT visualized across all other samples and replicates, ordered most to least frequent. Dot size reflects estimated percentage representation of each barcode within a sample.

A pool of barcoded SK-N-SH cells was split into biological replicates with comparable barcode diversity. Replicates were exposed to cisplatin for seven days (T1) and subsequently cultured in drug-free medium for one week (T2) or four weeks (T3).

Single-cell transcriptomic analysis of untreated SK-N-SH cells identified canonical ADRN and MES cells as previously reported. A population of cells expressing mixed ADRN and MES features was also observed, hereon referred to as bridge (BRI) cells (Figure 1B, Figure S1A-B, Table S1 and S7). To better understand the dynamics between these cell states we interrogated the scRNA-seq data using pseudotime analysis. We observed that ADRN, BRI and MES cells reside in a phenotypic continuum with BRI cells occupying a transitioning position in ADRN-to-MES trajectories (Figure S1A). To further characterise cell state transitions in neuroblastoma, we modelled them as a multiscale switch process using dynamical manifold approaches (MuTrans)^45^, an alternative to the entirely continuous perspective presumed by pseudotime analysis. This further confirmed that neuroblastoma cell states reside within a phenotypic landscape composed of semi-stable identities connected by transitional paths (Figure 1C). Low entropy states (i.e. committed) were heavily composed of MES or ADRN cells, while high entropy wells (i.e. plastic) contained the highest level of BRI cells (Figure S1B-C). Moreover, the most probable transitional paths between MES and ADRN wells involved conversions through BRI states, highlighting the role of these cells as mediators of plasticity (Figure 1C). Functional isolation of BRI cells using CD44-based FACS^29^ revealed their enhanced colony-forming ability and phenotypic instability (Figure S1D-F).

To study the role of neuroblastoma’s innate plastic potential in adaptation to treatment, we monitored phenotypic dynamics across the cisplatin treatment and recovery time course. After cisplatin treatment (T1), the original phenotypic heterogeneity was eradicated, with the surviving population exhibiting a predominant MES identity. These cells also expressed markers of senescence, compatible with entrance into a drug-tolerant persister (DTP)-like state (Figure S2). Upon cisplatin withdrawal, a gradual re-emergence of phenotypic heterogeneity was observed (T2, awakening), which culminated into a total recovery of the original proliferative capacity and phenotypic composition (T3) (Figure 1B). GSEA of the genes specifically expressed in T3 vs T2 revealed that once recovered, neuroblastoma cells downregulate plasticity (EMT) and proliferation programmes (Figure S2G).

We next wanted to know whether the adaptation process, comprising selection of MES-DTPs followed by plasticity-led awakening, was mediated by pre-existing lineages. To unravel the contributions of lineage selection and phenotypic transitions, we leveraged the expressed DNA barcodes to trace clonal dynamics throughout the treatment response framework.

Plasticity played a minimal role in the initial phase of adaptation, yet it was essential for the second adaptation phase, involving awakening and regrowth through MES to ADRN transitions. Lineage tracing analysis using expressed barcodes and single-cell transcriptomic information revealed that awakened lineages (those that survive treatment and are present at T3) were already composed of a majority of MES cells before treatment (Figure 1D and S3D). At persistence, we observed a selection of these pre-existing MES lineages, rather than ADRN-to-MES plasticity. During awakening after drug removal, some of these lineages remained highly MES, while others underwent plasticity into ADRN and BRI identities. To account for the limitations of this approach, we also calculated the frequency of cells in each cell identity for the barcode shared across all timepoints, for which we observed comparable dynamics (Figure S1E). Further functional approaches showed only pure MES populations – and not pure ADRN ones – were able to recover from cisplatin treatment, through plasticity towards ADRN states (Figure S3E). Altogether, these results strongly support treatment selects for phenotypes and plasticity emerges post treatment.

To test the contribution of plasticity to each phase of adaptation as a falsifiable hypothesis, we applied the mathematical framework described by Whiting et al.^46^ to the lineage information derived from our DNA barcodes. Their model predicts that when phenotypic changes at the population level are mediated by selection of pre-existing fit clones, the same lineages are repeatedly selected across independent replicates, leading to low inter-replicate barcode diversity. Conversely, when phenotypic plasticity operates, a different set of lineages can transition and dominate in each replicate, generating high inter-replicate barcode diversity. At T1, we observe very limited inter-replicate diversity (1.25; theoretical range 1-3; Figure 1F and S3F) and limited changes in frequency distributions of barcodes (Figure S3A-C). Despite the limitations in resolution of the barcoding system, this suggests that the MES-DTP enrichment observed during early adaptation is driven not by selection at the level of lineages but cell-states. In contrast, upon awakening and regrowth (T3) inter-replicate barcode diversity increases sharply to 2.14 (Figure 1F and S3F). This is consistent with a plasticity-driven stochastic process in which only a small, replicate-specific subset of clones contributes disproportionately to regrowth.

### AP-1 activity increases post-treatment and promotes plasticity-led awakening and recovery

We next aimed to identify the molecular determinants underlying this two-step adaptive strategy to gain insights into the earliest stages of treatment resistance. To this end, we employed single-nucleus multiomics in a sample of barcoded untreated and recovered SK-N-SH cells, enabling simultaneous profiling of chromatin accessibility (snATAC-seq) and gene expression (snRNA-seq) within the same cell (Table S1).

Transcriptomic characterisation of the cells revealed a comparable MES-BRI-ADRN continuum to the observed in scRNA-seq data. Moreover, transfer of this annotation to the matched ATAC-seq profiles, showed that cells of the same cell identity displayed similar chromatin landscapes (Figure 2A-B).

**Figure 2.**
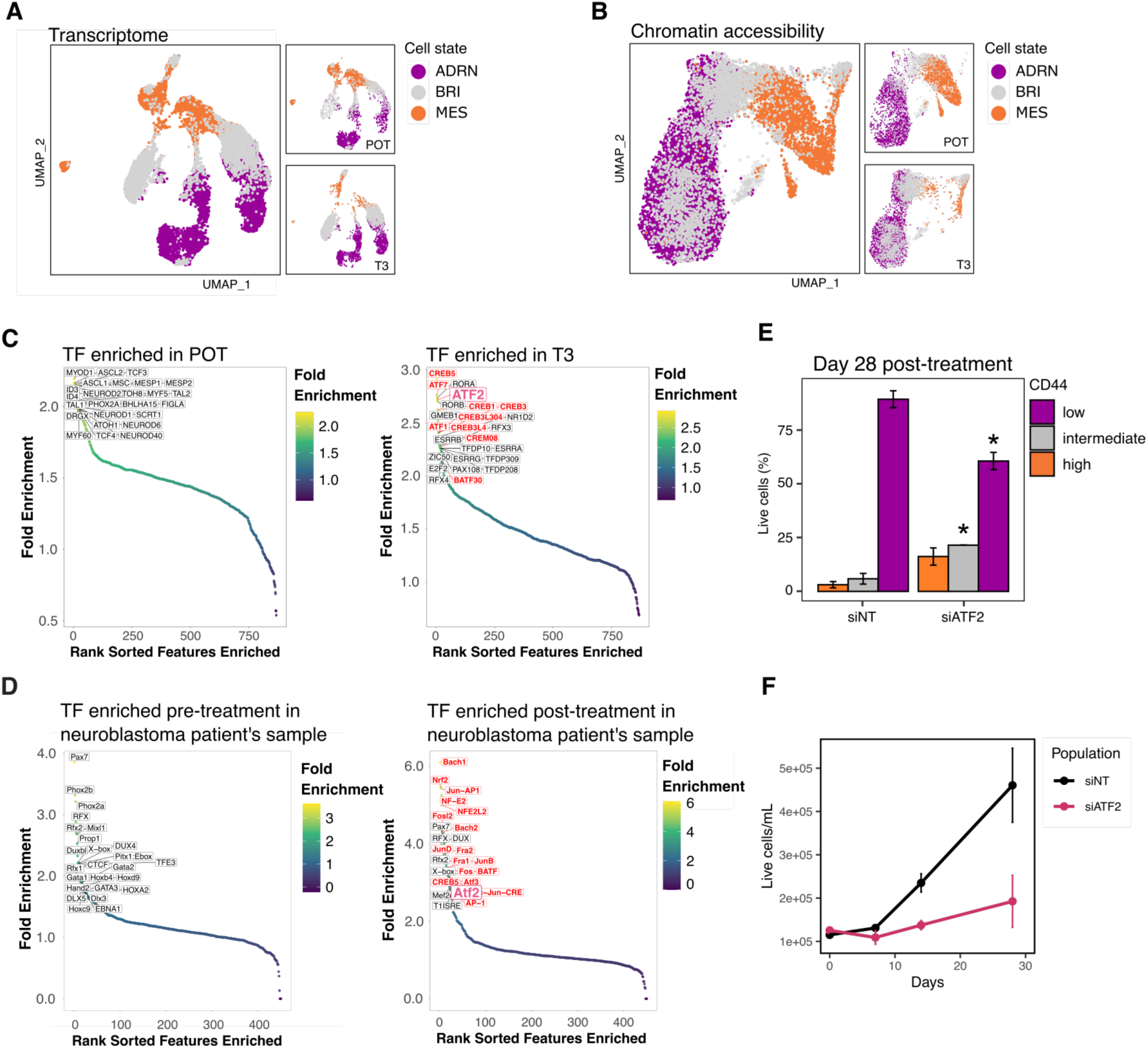
AP-1-mediated chromatin rewiring enables plasticity-led adaptation post-treatment. (A) Umap based on transcriptomic (A) or chromatin accessibility (B) information in untreated (POT) and recovered after cisplatin treatment (T3) SK-N-SH cells. Cells were annotated into ADRN, BRI or MES using the transcriptomic information (A), as described in Methods. This annotation was then transferred to the ATAC-seq profiles (B) based on matched nucleus ID. (C) Top 15 transcription factor binding motifs enriched in the differentially accessible peaks in ADRN cells in POT (left) or T3 (right) populations. Transcription factors highlighted in red are members of the AP-1/bZIP family. (D) Top 25 transcription factor binding motifs enriched in the differentially accessible peaks in ADRN-Calcium cells at diagnosis (left) or after treatment (right) in 22 high-risk neuroblastoma patients as described in *Yu et al.*^47^. (E) Percentage of CD44-high (MES), intermediate (BRI) and low (ADRN) cells in SK-N-SH cells upon cisplatin treatment and down-regulation of ATF2. Cells were treated with an IC50 concentration of cisplatin for 7 days, after which they were transfected with a siRNA against ATF2 (siATF2) or a non-targeted control (siNT). CD44 expression was assessed by flow cytometry 28 days after cisplatin treatment. Bars represent the average of three replicates. Statistical significance is denoted using asterisks (*P ≤ 0.05) based on t-test comparing the results to the siNT control. (F) Cell growth (measured as change in concentration of live cells) during recovery from cisplatin treatment upon knockdown of ATF2 in SK-N-SH cells. Dots represent the average of three replicates.

We observed that although transcriptomic profiles remained broadly comparable between untreated and recovery (T3) time points, chromatin accessibility landscapes were substantially altered at recovery, suggesting that cisplatin treatment induces sustained chromatin rewiring (Figure S3F-G, Table S2).

To identify regulators of plasticity-led awakening and regrowth, we focused on T3-ADRN cells, which had transitioned from MES-like DTPs, and compared them to ADRN cells from untreated conditions. Cells at recovery displayed significantly increased chromatin accessibility across the genome compared to their untreated counterparts (Figure S3H). Motif enrichment analysis revealed that the differentially accessible regions were strongly enriched for binding sites of early stress response transcription factors in the AP-1/bZIP family (Figure 2C).

To assess whether AP-1 activation also occurs in patient tumours, we analysed AP-1 activity patterns using longitudinal single-nucleus multiomics data of 22 high-risk neuroblastoma patients^47^. We found that cell identities that persist and tolerate induction therapy, particularly those associated with poor clinical outcome, displayed significantly increased chromatin accessibility at AP-1 binding motifs after treatment compared to diagnosis. (Figure 2D).

To functionally test the role of AP-1 in treatment adaptation, we focused on ATF2, a well-characterised member of the AP-1/bZIP family, and one of the top binding motifs significantly enriched in recovered populations in cell lines and patient samples in our previous analysis (Figure 2C-D). We used an siRNA approach to downregulate ATF2 in SK-N-SH cells and subjected them to the same ON/OFF cisplatin treatment framework (Figure S3I). Reduction of ATF2 levels had minimal effects on baseline cell viability, but upon drug withdrawal, siATF2 cell cultures exhibited markedly delayed recovery kinetics, reduced proliferative rebound, and impaired restoration of the phenotypic diversity normally regained during awakening (Figure 2E-F). Specifically, ATF2 depletion blocked MES-to-ADRN plasticity post-treatment, preventing the re-emergence of proliferating ADRN cells from DTP-MES.

Together, recovery from cisplatin was accompanied by increased chromatin accessibility at AP-1/bZIP motifs in both SK-N-SH cells and patient tumours, and ATF2 depletion delayed recovery and impaired the MES-to-ADRN transition underlying awakening.

### Transcriptional noise at cell-identity regulators drives stochastic awakening from MES-DTP states

While therapy-induced AP-1 activity primes the chromatin for adaptive rewiring, the stochastic nature of DTP awakening and phenotypic diversification remains mechanistically unclear. Transcriptional noise, defined as the stochastic fluctuations in gene expression, is a well-described source of phenotypic diversification in otherwise homogeneous populations^20,23,24,48^. Given the probabilistic behaviour observed in the lineage diversification during post-treatment regrowth (Figure 1F–G), we hypothesised that transcriptional noise contributes to plasticity-led DTP awakening in neuroblastoma.

To address this, we examined the timing and targets of transcriptional noise across the ON/OFF treatment framework in neuroblastoma preclinical models. We used the BASiCS statistical framework^49,50^, restricting our analysis to cells within the same phenotypic annotation to avoid confounding effects from cell state heterogeneity. This allowed us to assess noise dynamics within a specific cell identity as cells transitioned from persistence into awakening.

While noise levels remained comparable across timepoints in ADRN and BRI cells (Figure S4A-B), we observed a dynamic pattern in MES cells, which are the substrate for plasticity-led regrowth. In untreated MES populations, 2.7% of genes were classified as highly variable, consistent with neuroblastoma populations existing in phenotypic landscapes with some levels of innate plasticity (Figure 3A). This fraction decreased during persistence (1.3%) but rose sharply upon treatment withdrawal during awakening and regrowth (8.0%), when plasticity is at its maximum. Once the population recovered and heterogeneity was restored, transcriptional noise levels dropped back to baseline (3.0%) (Figure 3A-B; Table S3). Importantly, transcriptional noise fluctuations were not correlated with cell-cycle state or viability, indicating that changes in noise reflect regulatory variability rather than proliferative differences (Figure S4C-D). Moreover, inference of changes in transcriptional bursting parameters^51^ between persistence and awakening revealed an enrichment of noise-causing bursting patterns in highly variable genes, (Figure S4E-G) highlighting the intrinsic origin of the transcriptional noise dynamics observed. These results establish that transcriptional noise and plasticity are tightly coupled during the drug response process.

**Figure 3.**
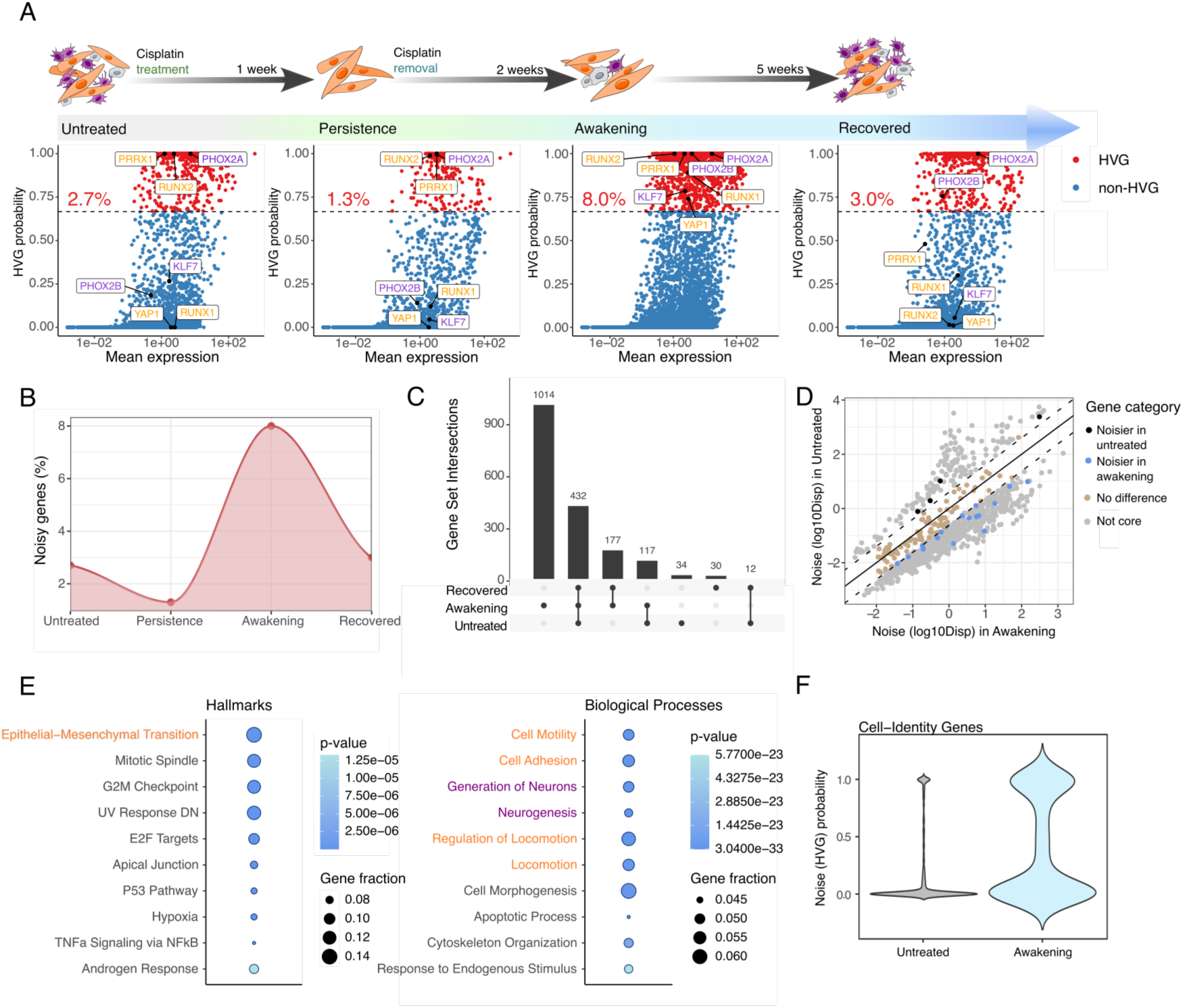
Plasticity-led awakening from MES-DTPs is enabled by transcriptional noise in cell identity programs. (A) High Variable Genes (HVG) analysis in MES cells across stages of cisplatin treatment. Dots represent individual genes, dashed lines statistical threshold and numbers percentage of the whole transcriptome that corresponds to HVG genes. Boxes label important MES (orange) and ADRN (purple) master regulators. (B) Percentage of noisy genes across treatment stages. (C) Overlap of high noise genes across stages of cisplatin treatment. (D) Differentially noisy genes between untreated and awakening MES cells (grey dots). Coloured dots highlight genes included in the core of noisy genes (shared between untreated, awakening and recovered): black, higher noise in untreated; blue, higher noise in awakening; brown, no difference between timepoints. (E) GSEA of high noise genes specific to the awakening timepoint. Highlighted text relates to cell-state identity pathways in MES and ADRN cells. (F) Noise probability of MES and ADRN identity genes (*van Groningen et al.*^40^) in untreated and awakening MES cells.

To identify genes underlying this phenomenon, we characterised the noisy gene pool across treatment stages. We uncovered a core set of 432 genes that remained highly variable before and after treatment (Figure 3C). During awakening, while the genes in the core remained equally variable or gained extra noise, transcriptional noise expanded to include hundreds of additional genes (Figure 3C-D). Awakening-specific genes were enriched for plasticity-associated processes (e.g. EMT) and cell identity programs (e.g. cell adhesion, motility, neurogenesis) (Figure 3E), including important regulators of the ADRN and MES identities, namely, PHOX2A, PHOX2B, KLF7 and PRRX1, RUNX1, RUNX2, YAP1, respectively^40,41^ (Figure 3A, E-F). Notably, these genes exhibited minimal changes in mean expression and no consistent directionality, underscoring that transcriptional noise is the primary driver of expression variability (Figure S4H-J).

Finally, to establish a causal relation between transcriptional noise and the emergence of plasticity, we treated neuroblastoma models with the base analogue and noise enhancer 5-iodo-2’-desoxiuridine (IdU)^20^. Treatment with IdU induced transcriptional noise in SK-N-SH cells, measured as the increase in expression variability of a destabilised GFP (d2GFP) reporter^20^ (Figure S5A-B). This resulted in an increased migratory potential and enrichment of BRI cells (Figure S5C-F). Induction of noise also increased the migratory features and the activation of plasticity-related expression programs in a patient-derived model cultured *ex vivo* (Figure S6).

### Transcriptional noise dynamics during therapy response are conserved across neuroblastoma patient-derived models and patient samples

We then sought to evaluate whether the link between transcriptional noise and plasticity-led recovery observed in preclinical cell lines extends to patient-derived neuroblastoma models. To this end, we analysed scRNA-seq data of two neuroblastoma patient-derived organoid models (PDOs; NB039 and NB067) and a patient-derived xenograft (PDX; GR-NB5) cultured *ex vivo*. Despite model-specific variation in the gene sets defining cellular states across models (Figure S7A-H), treatment-induced phenotypic dynamics were highly conserved (Figure S7I-J). To facilitate cross-model comparisons, we annotated cell states based on their dynamics in response to treatment rather than by gene identity (Figure S7I). Cisplatin reduced highly proliferating cells (PROL) and increased treatment-resistant reservoir cells (RES – akin to the MES phenotype in pre-clinical cell lines). Upon drug withdrawal, the awakening phase was characterised by the enrichment of previously rare bridge cells (BRI), marking phenotypic transitions towards repopulation and restoration of heterogeneity. Consistent with this functional annotation, BRI cells exhibited the highest transcriptional noise levels, further supporting their role as biomarkers of plasticity (Figure S8A).

To investigate transcriptional noise dynamics within the dominant DTP phenotype, we applied BASiCS to DTP-RES cells at different treatment phases (Figure 4A and S8B; Table S4). In both PDO models, transcriptional noise increased markedly during the awakening phase preceding recovery, with 4.5% (NB039) and 4.9% (NB067) of genes classified as highly variable, compared with 2.4% and 1.5%, respectively, in the pre-treatment condition. In contrast, DTP-RES cells from the GR-NB5 model did not display significant changes in transcriptional noise, consistent with the absence of recovery in this model upon drug withdrawal (Figure 4A and S7J).

**Figure 4.**
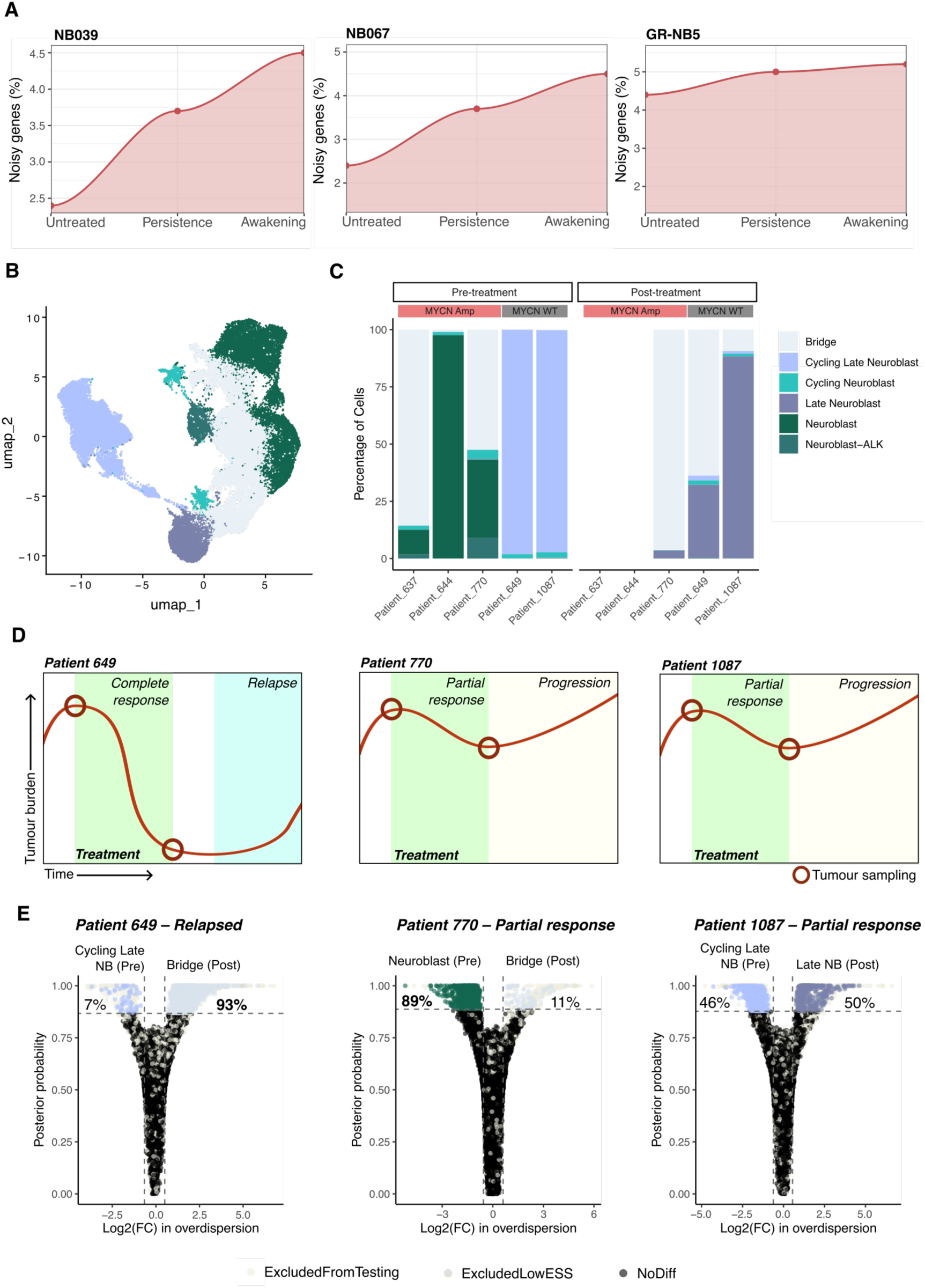
Transcriptional noise increases after treatment and precedes relapse in patient-derived models and samples. (A) Percentage of noisy genes across treatment stages in three patient-derived models cultured *in vitro*. (B) Umap showing phenotypic heterogeneity in pre-and post-treatment samples of five neuroblastoma patients. Transcriptomic clusters were annotated based on the expression of canonical markers of the developing sympatho-adrenal lineage^39^. (C) Phenotypic composition across patients and treatment stages. (D) Patient clinical history and sampling times. (E) Differential noise analysis between the dominant population pre-and post-treatment in matched patient samples. Percentages represent the fraction of noisy genes that display higher variability in each identity/timepoint.

Consistent with our findings in cell lines, we identified a core gene set that maintains high noise levels before and after treatment, encompassing genes involved in cell-cycle progression, translation, and neuronal phenotype regulation. During awakening, this noisy gene pool expanded substantially, incorporating a broader cohort of genes associated with neuroblastoma cell identity programmes (Figure S8C-D)

To interrogate the translational implications of our discoveries, we asked whether transcriptional noise dynamics could also be observed in longitudinal samples from neuroblastoma patients. We performed single-nuclei RNA sequencing on flash-frozen neuroblastoma tumour samples, including three matched pre-and post-treatment samples from patients receiving a combination of etoposide, carboplatin and vincristine (Table S5). Malignant cells harbouring typical neuroblastoma chromosomal aberrations were annotated into cell states based on the expression of established sympatho-adrenal developmental markers^39^ (Figure S9). At diagnosis, tumours segregated by MYCN status (Fig. 4B–C), with MYCN-amplified samples enriched for neuroblast and progenitor-like (bridge) states, whereas MYCN wild-type tumours were predominantly composed of more differentiated cycling late neuroblasts.

Consistent with our preclinical observations, treatment altered the cell state composition of each patient’s tumour. Across all matched pre-and post-treatment samples, we observed a reduction in proliferative and more differentiated-like cell populations, accompanied by a relative enrichment of progenitor-like states (Figure 4C).

To examine the relationship between cell-state composition, transcriptional noise and clinical outcome, we performed a differential noise analysis for each patient by comparing the dominant cell identity before and after treatment (Figure 4D-E). Patient 649 displayed a drug response trajectory closely resembling that of our dynamic preclinical models: an initial response followed by the awakening of residual DTPs and eventual relapse. In this patient, we observed a marked increase in transcriptional noise within cell states that had persisted through treatment, consistent with our preclinical observations. In contrast, patients 770 and 1087 exhibited intrinsic resistance, showing only partial responses that progressed to treatment failure. These tumours were not suitable comparators for our preclinical awakening model and exhibited either elevated pre-treatment noise or no significant change in noise across time points, respectively.

Collectively, an increase in transcriptional noise after treatment was observed in the patient whose clinical course matched the response-awakening-relapse trajectory, but not in the two patients with intrinsic resistance.

### KDM5 inhibition reduces transcriptional noise and blocks awakening and regrowth

Our results thus far demonstrate that plasticity is necessary for DTP exit and that this process is underpinned by transcriptional noise, identifying noise as a potential evolutionary vulnerability in cancer. However, targeting a probabilistic process such as transcriptional noise presents a fundamental challenge. To address this, we turned to chromatin regulators known to influence gene expression variability, a readout of transcriptional noise, through chromatin rewiring. We focused on the KDM5 family of histone demethylases, which remove H3K4me3 marks from gene promoters for two reasons: (1) KDM5B is a core component of the MES signature in neuroblastoma^40^, suggesting its functional relevance in maintaining the MES cell state identity; and (2) KDM5 inhibition via the compound KDM5-C70 was shown to reduce gene expression variability in breast cancer^52^. To investigate its role in noise-driven adaptive evolution in neuroblastoma, we first explored its expression level during the dynamic drug response process. We detected an increase in KDM5A/B protein expression upon cisplatin withdrawal, suggesting that KDM5 activity is dynamically upregulated during recovery and may facilitate the epigenetic reprogramming that drives transcriptional noise and enables plasticity-led awakening post-treatment (Figure S10A)

To further test the hypothesis that KDM5 regulates transcriptional noise and influences plasticity-led recovery, we experimentally modulated KDM5 activity and assessed its impact on gene expression variability. Using SK-N-SH cells expressing a destabilised GFP (d2GFP) reporter, we measured transcriptional noise following treatment with IdU (a known noise inducer), cisplatin, and the KDM5 inhibitor KDM5-C70. IdU significantly increased expression variability as expected^20^, while cisplatin treatment had a limited effect. Remarkably, KDM5 inhibition led to a modest but significant reduction in transcriptional noise (Figure S10G). Similarly, a subset of genes that displayed high noise levels during awakening reduced their noise probability upon KDM5 inhibition, including markers of neuroblastoma cell identity (Figure S10H-I).

We next tested whether KDM5 regulation of transcriptional noise functionally affects plasticity-led awakening and regrowth following treatment. To do this, we applied our preclinical ON/OFF treatment framework encompassing the persistence, awakening, and recovery phases (Figure 1A), and administered KDM5-C70 either as a single dose or in weekly intervals following cisplatin withdrawal. Compared to the control arm, KDM5 inhibition markedly delayed MES-DTP exit through MES-to-ADRN transitions, with weekly dosing producing the strongest effect (Figure 5B). Pre-treatment with KDM5-C70 before cisplatin withdrawal produced a similar outcome (Figure S10B). Importantly, in untreated SK-N-SH cells, KDM5-C70 showed minimal toxicity and did not alter phenotypic composition, excluding a confounding factor due to selective cytotoxicity (Figure S10C–D). Knock-down of KDM5A had a similar effect to chemical inhibition, although not to the same extent, possibly due to residual KDM5A activity and overlapping functions with other KDM5 family members, including KDM5B (Figure S10E-F).

**Figure 5.**
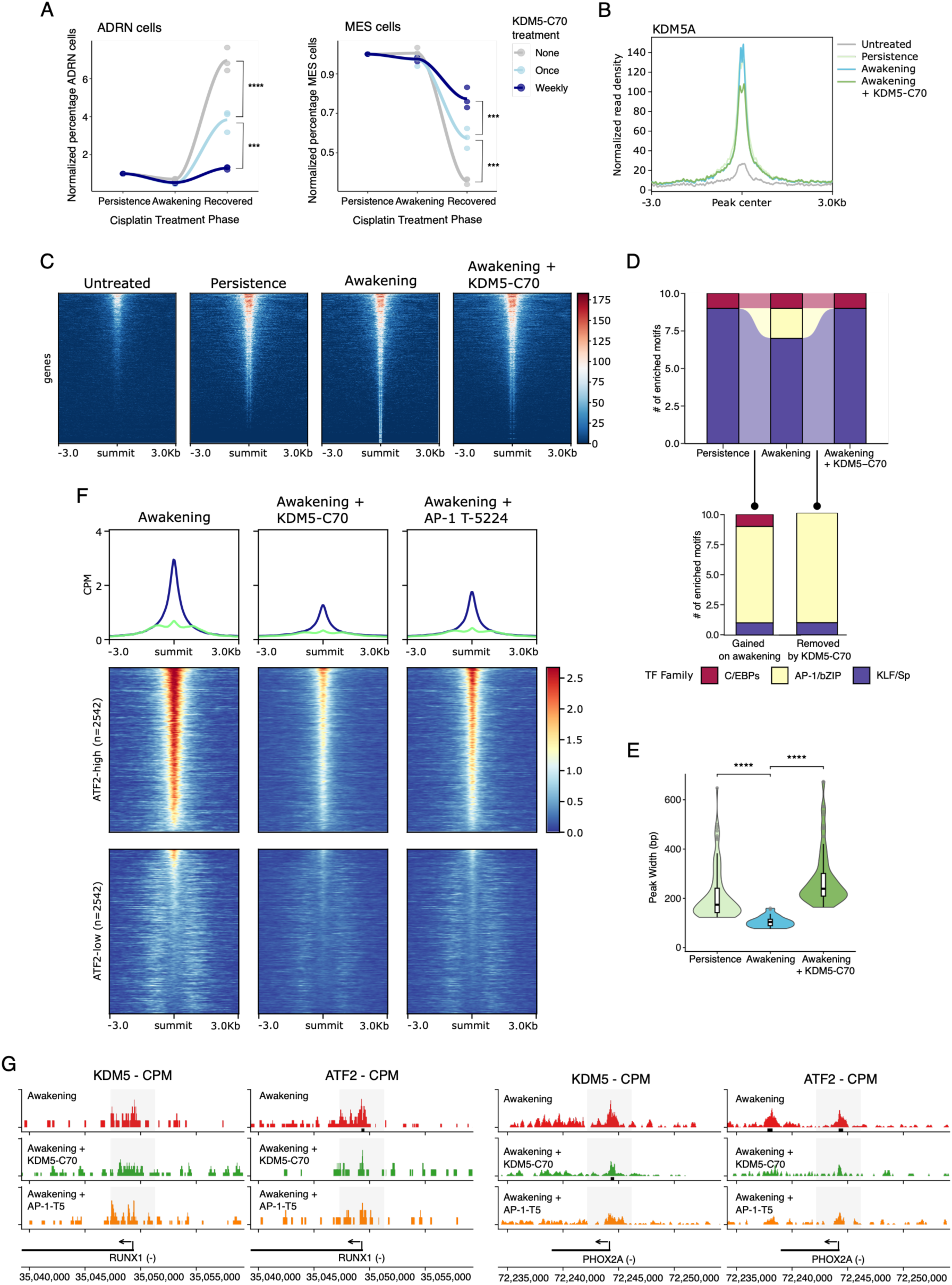
KDM5A binding increases during awakening and is guided by AP-1 to cell identity loci. (A) Change in ADRN and MES cells during stages of cisplatin treatment in the absence of the KDM5 inhibitor KDM5-C70 or when it is added only once or weekly after cisplatin removal. Data points were normalized to the percentage of each phenotype on the day of cisplatin removal. Stars represent statistical significance resulting from a one-way ANOVA test with Tukey’s HSD (‘****’ p <0.0001, ‘***’ p <0.001, ‘**’ p<0.01, ‘*’ p <0.05, ‘ns’ p >0.05’). (B, C) Genome-wide KDM5A chromatin occupancy across stages of cisplatin treatment and upon KDM5 inhibition. (B) Cumulative read density across KDM5A peaks differentially bound between the untreated and each of the treated conditions. (C) Signal intensity of KDM5A binding to differentially bound peaks between untreated and the remaining conditions. Each panel contains all peaks differentially bound across all conditions vs. untreated. Peaks are ordered and coloured by signal intensity. (D) Characterisation of the top 10 most enriched transcription factor binding motifs in the proximity of differential-KDM5 peaks across stages of cisplatin treatment and upon KDM5 inhibition. (E) Width of H3K4me3 peaks in noisy genes during awakening across cisplatin treatment stages. Stars represent results of a Wilcoxon rank sum test (‘****’ p <0.0001). (F) Counts per million (CPM) in KDM5A peaks across conditions. Mid and bottom panels display signal intensity of KDM5A peaks, ordered from highest to lowers intensity. (G) Track plots of KDM5A and ATF2 occupancy at RUNX1 and PHOX2A during awakening in the presence or absence of KDM5 or AP-1 inhibition.

KDM5 inhibition therefore reduced transcriptional noise and delayed MES-to-ADRN transitions following cisplatin withdrawal, without detectable cytotoxicity or effects on phenotypic composition in untreated cells.

### AP-1 primes KDM5 chromatin rewiring to enhance transcriptional noise at cell identity genes

To dissect the mechanism by which KDM5 activity regulates plasticity-led awakening and regrowth, we analysed the chromatin-binding dynamics of KDM5A, KDM5B, and H3K4me3 across the established preclinical drug-response trajectory. KDM5B binding remained low and largely unchanged throughout the drug-response phases (Figure S11A). In sharp contrast, KDM5A binding increased following cisplatin exposure. KDM5A signal increased in 23% of all peaks detected in untreated cells, additionally 2,840 new KDM5A peaks were detected. This remodelling of the KDM5A-biding landscape intensified during awakening, with 26% of the untreated peaks increasing in signal and 8,068 new peaks gained. Nevertheless, this was significantly reduced by KDM5-C70 treatment, leading to only 6,189 new peaks being gained from the untreated condition (Figure 5C-D).

When we focused specifically on neuroblastoma cell identity genes and noisy genes we had previously identified as being linked to the plasticity-led awakening process, we observed similar patterns of increased KDM5A binding post-treatment and inhibition of this process by KDM5-C70 (Figure 5E-F and S11B-C).

Transcription factor motif analysis of the awakening specific peaks revealed a significant over-representation of AP-1 binding motifs, especially at enhancers, implicating AP-1 in guiding KDM5A chromatin recruitment (Figure 5G and S11D). This is consistent with our earlier findings showing AP-1 sites become more accessible following treatment, and that ATF2 downregulation impairs post-treatment phenotypic recovery. Taken together, these results suggest that AP-1 acts upstream to remodel chromatin and prime KDM5A engagement at identity loci, thereby facilitating transcriptional noise-mediated plasticity.

We next examined whether these binding dynamics influenced H3K4me3, the canonical substrate of KDM5A. Although global H3K4me3 levels rose during persistence, they remained largely unchanged during awakening or upon KDM5 inhibition (Figure 5H and S11E). However, we observed a significant decrease in H3K4me3 peak width at promoters of noisy genes during awakening (median 104 bp versus 174 bp at persistence). This effect was rescued by KDM5 inhibition (239 bp; Figure 5I). The narrowing of H3K4me3 peaks, rather than global demethylation has been previously associated with increased transcriptomic heterogeneity in cancer cell populations^52^.

Overall, KDM5A binding increased at cell-identity and noisy genes during awakening and was reduced by KDM5 inhibition. These awakening-specific peaks were enriched for AP-1 motifs, and H3K4me3 peaks at noisy gene promoters narrowed during awakening.

To understand the interplay between KDM5A and AP-1in the chromatin rewiring that drives plasticity-led awakening, we profiled the chromatin binding landscape of KDM5A and its substrate H3K4me3 together with ATF2, c-Jun and c-Fos, three AP-1 family members that commonly co-occupy AP-1-responsive regulatory elements^53,54^. We identified genomic regions enriched for H3K4me3 and stratified them by ATF2 occupancy. KDM5A binding was consistently enriched in regions also bound by ATF2, and by c-Jun and c-Fos (Figure 5F and S12). Significantly, pharmacological inhibition of AP-1 activity with T-5524^55^ reduced KDM5A binding to levels comparable to direct KDM5 inhibition (Figure 5F and S12), and decreased noise probability at highly variable genes during awakening (Figure S12E). The same pattern was observed at cell-state identity genes that gain noise during awakening (Figure S12A-D; see also Figure 3A).

These results indicate that KDM5A and AP-1 factors co-occupy regulatory regions, including those of noisy cell-identity genes, during awakening, and that AP-1 activity is required for KDM5A recruitment to these loci.

### Transcriptional noise drives phenotypic plasticity-led adaptation to treatment in hepatoblastoma models

Finally, we wanted to investigate whether transcriptional noise similarly contributes to plasticity-led recovery in other paediatric cancers with low genomic instability. We focused on hepatoblastoma, the most common paediatric liver malignancy and a tumour type characterised by one of the lowest mutational burdens among paediatric cancers^4^. Emerging evidence, including a parallel study from our group, indicates that phenotypic plasticity plays a critical role in hepatoblastoma adaptation to treatment^30,56^. Using DNA-barcoded lineage tracing and single-cell multiomics in HuH6 and HepG2 hepatoblastoma cell lines, patient-derived xenografts and organoids, and longitudinal patient samples, our work shows that cisplatin selects a progenitor-like drug-tolerant persister state, from which post-treatment regrowth proceeds through plasticity-driven re-diversification along a phenotypic continuum spanning liver developmental stages, a two-step adaptive strategy mirroring our observations in neuroblastoma^57^.

Hepatoblastoma cell populations exist in a phenotypic continuum, similar to neuroblastoma, along phenotypic landscapes whose semi-stable states resemble different stages of liver development^57^. Specifically, HuH6 included cell states with features of slow-proliferating, undifferentiated progenitor-like cells and more differentiated, proliferative states resembling hepatocytes. These states follow similar dynamics during treatment to those observed in neuroblastoma, and thus we have relabelled them for consistency across the two datasets. Hepatocytic-like proliferative states (hereafter PROL-I and PROL-II) dominated the untreated populations and were especially sensitive to treatment; progenitor-like states separated into those that dominated during persistence and occupied extreme UMAP positions (hereafter RES) and those that emerged during awakening and had intermediate positions (hereafter BRI)^57^ (Figure S14A-D).

Mirroring our observations in neuroblastoma, plasticity drives the awakening and regrowth of RES-DTP cells post cisplatin treatment, leading to a recovered population through phenotypic transitions from progenitor-like to differentiated-like states, restoring the originating phenotypic heterogeneity^57^ (Figure S14D).

To investigate the role of transcriptional noise in hepatoblastoma adaptation, we quantified noise levels in RES cells across different stages of the treatment response framework. This analysis revealed dynamic changes in transcriptional noise that closely mirrored those observed in neuroblastoma models, with noise peaking during the post-treatment awakening phase. The highest levels of transcriptional noise were observed during awakening, with levels of highly variable genes resetting to pre-treatment values after recovery (Fig. 6A-B; Table S6). As we had uncovered in neuroblastoma, we observed a core of 152 noisy genes at baseline, which, during awakening expanded to include thousands of additional genes, primarily involved in cell cycle, differentiation and plasticity regulation. These included genes known to drive plasticity during cisplatin treatment in hepatoblastoma, as well as markers of RES and PROL cells^57^ (Figure 6A and S14E-F). Noise dynamics in hepatoblastoma HuH6 cells therefore mirrored those in neuroblastoma SK-N-SH cells.

**Figure 6.**
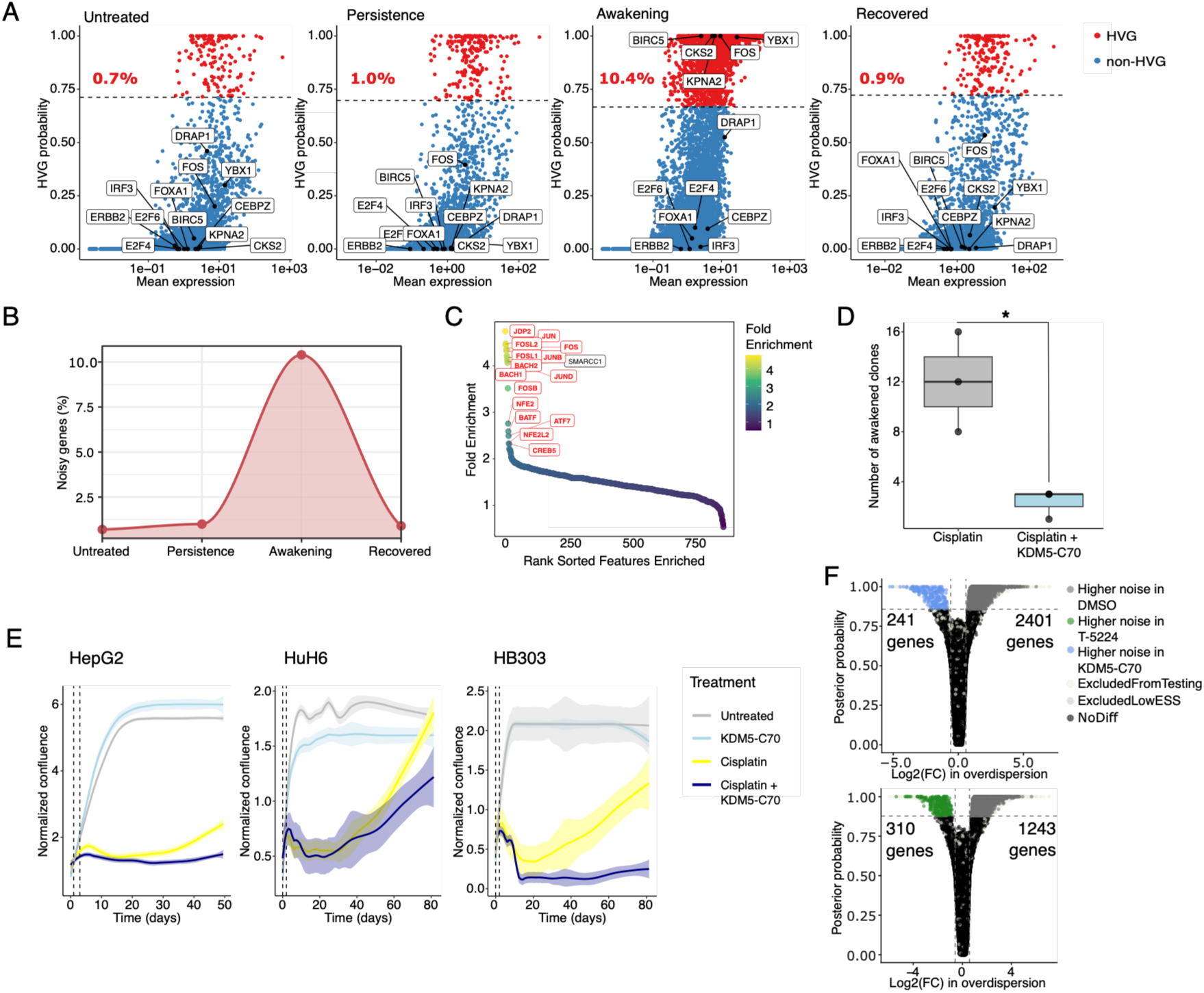
KDM5-mediated transcriptional noise regulates plasticity-led adaptation in hepatoblastoma models. (A) High Variable Genes (HVG) analysis in RES (DTP-like) cells across stages of cisplatin treatment in the hepatoblastoma cell line HuH6. Dots represent individual genes, dashed lines statistical threshold and numbers percentage of the whole transcriptome that corresponds to HVG genes. Boxed highlight genes involved in plasticity-led treatment adaptation. (B) Percentage of noisy genes across treatment stages. (C) Top 15 transcription factor binding motifs enriched in differentially accessible peaks HuH6 cells after recovery from cisplatin treatment, compared with untreated cells. (D) Number of clones after awakening in HepG2 hepatoblastoma cells treated with cisplatin for seven days and KDM5-C70 once after cisplatin removal. Stars represent statistical significance resulting from a t-test (‘****’ p <0.0001, ‘***’ p <0.001, ‘**’ p<0.01, ‘*’ p <0.05, ‘ns’ p >0.05’). (E) Confluency of HepG2, HuH6 and HB303 hepatoblastoma cells treated with cisplatin for two days. Lines represent average of two or three replicates; shaded areas represent standard deviation of the mean. Dashed lines highlight day of cisplatin treatment and removal. Confluency was normalized to the percentage on the day of cisplatin removal. (F) Volcano plots displaying pairwise comparisons of transcriptional noise levels (overdispersion) in awakening HuH6 in the presence or absence of KDM5A (KDM5-C70) or AP-1 inhibition (T-5224). Each dot represents a gene and are coloured by the condition at which they are significantly noisier.

We next analysed the HepG2 hepatoblastoma cell line, which, in contrast to HuH6, exhibited low transcriptional noise across all cell states and treatment phases, with no significant dynamic changes observed (Figure 15A-D). We speculate that this is due to a rapid recovery of this model, which is dominated by hepatocytic cells by the awakening sampling timepoint, such that the real awakening window was missed at our timepoint (Figure S15C). Nonetheless, despite the overall lower noise levels, we identified a shared subset of noisy genes between HuH6 and HepG2, enriched in overlapping biological processes (Figure S15E-F).

We then asked whether, as observed in neuroblastoma, the AP-1/KDM5 axis might regulate noise levels during treatment response in hepatoblastoma. To test this, we analysed single-nuclei multiomics data from untreated and recovered (T3) HuH6 cells^57^. We observed that AP-1 binding sites were amongst the most accessible sites at recovery (Figure 6C).

To functionally test the contribution of KDM5 and AP-1 to awakening, we treated HuH6, HepG2 and the patient-derived hepatoblastoma model HB303 with KDM5-C70 following cisplatin treatment. KDM5 inhibition halted awakening of DTP-RES HepG2 and HB303 cells and moderately delayed this process in HuH6 cells (Figure 6D-E). We further inhibited KDM5 or AP-1 activity by treating HuH6 cells with KDM5-C70 and T-5224. Analysis of transcriptional noise as performed above (Figure 3) on snRNA-seq showed that inhibition of either KDM5A or AP-1 led to a significant reduction in transcriptional noise during awakening (Figure 6F).

Overall, transcriptional noise in hepatoblastoma peaked during awakening mirroring our observations in neuroblastoma, AP-1 motifs were among the most accessible sites at recovery, and KDM5 or AP-1 inhibition reduced transcriptional noise and delayed or prevented awakening across hepatoblastoma models.

Taken together, our findings support a model in which transcriptional noise acts as a regulated driver of diversification that precedes and fuels phenotypic evolution in paediatric cancers. This process is mediated by AP-1 and KDM5 chromatin remodelling, and specifically targets genes associated with cell state identity and their regulation. Whether this mechanism is conserved across other paediatric and adult cancers remains an open question for future investigation.

## DISCUSSION

Childhood cancers are characterised by disruptions in developmental processes^2,61^, allowing tumours to exploit embryonic mechanisms such as phenotypic plasticity to adapt without requiring additional genetic mutations. However, direct evidence linking plasticity to cancer evolution has been limited due to technological constraints.

Here, through the integration of expressed DNA barcoding, single-cell multi-omics, dynamic manifold modelling, and chromatin accessibility and binding landscape profiling, we provide a framework that resolves this long-standing question. We show that plasticity, rather than genetic selection, drives early adaptive evolution in paediatric tumours, allowing them to acquire high-fitness phenotypes without new mutations.

A substantial fraction of cancers are thought to evade treatment through non-genetic mechanisms of adaptation, often involving entry into and exit from drug-tolerant persister (DTP) states with quiescent features^62^. This process may be particularly relevant to paediatric cancers, whose genomic silence limits the availability of mutational routes for adaptation^4^. In previous studies in neuroblastoma, the most common extracranial solid tumour in children, have linked phenotypic transitions between the two prototypical cell states ADRN and MES, with invasiveness and treatment resistance^40–42^. However, whether such phenotypic dynamics arise through selection of pre-existing subpopulations or true cell state interconversion has been unclear due to technological constraints.

In this work, we show that transcriptional noise, a key biological mechanism of living cells that leads to stochastic variability in gene expression, is dynamically regulated in response to treatment to drive the phenotypic changes that underlie adaptive evolution. Through functional approaches and mathematical modelling of single-cell transcriptomic data, we show that paediatric cancer cell populations exist in phenotypic landscapes of model-specific topologies (see also Chen, English et al.^57^). In steady-state conditions, cells occupy semi-stable states of low variability in gene expression (low transcriptomic entropy). In neuroblastoma models, these correspond to the previously identified ADRN and MES states^40,41^. Continuous transitions through these states occur primarily through an intermediate cellular state, which we termed BRI cells. BRI exhibited high transcriptional entropy and showed phenotypic instability when isolated by flow sorting (Figure 1C and S1C-D), leading us to postulate that BRI cells are a readout of plasticity.

Following a strong bottleneck imposed by the chemotherapeutic agent cisplatin, in which populations were enriched for MES states, cells awakened and generated recovered populations that recapitulated the phenotypic heterogeneity of the pre-treated population, indicating entry into and exit from DTP states in our experimental setting. Significantly, BRI cells were enriched after treatment during awakening (Figure 1B), supporting plasticity peaking during this stage.

Lineage tracing using expressed static barcodes and functional analyses complemented these analyses, leading to our proposed two-step model for the early stages of adaptive evolution (Figure 1). The first step is primarily dependent on phenotypic selection, and the second step on probabilistic processes. Upon exposure to chemotherapy, phenotypic selection enriches for pre-existing MES lineages with the capacity to enter DTP states (MES-DTPs). Tumour regrowth after treatment withdrawal is instead not driven by a major selection process. Dormant MES-DTPs stochastically transition back to proliferative ADRN states, re-establishing phenotypic heterogeneity. We note, however, that barcode-based lineage tracing resolves clonal dynamics only above the detection threshold set by sequencing depth; selection acting on rare lineages can’t be excluded. Our data therefore argue against clonal selection as the dominant route to persistence and awakening, rather than excluding a contribution to it. Similar dynamics of dormancy followed by stochastic reactivation have been observed in adult cancers such as breast cancer, supporting the idea that plasticity-first evolution is a general principle of tumour relapse^63^. This two-step process fits within a broader framework of tumour evolution under fluctuating treatment pressure, in which selection can act not only on genetic clones^64,65^, but also on heritable, non-genetic states^66,67^.

Cytotoxic chemotherapy imposes a strong, transient selective pressure that eliminates sensitive cells and enriches for the small subset of tolerant cells, regardless of how tolerance is encoded^68^. Once treatment is withdrawn, this pressure relaxes – a phenomenon central to adaptive-therapy frameworks in which a flatter fitness landscape permits greater phenotypic exploration^69–71^. In our model, this relaxation of selection allows plasticity to dominate during awakening, without a deterministic pre-existing advantage dictating which lineage re-emerges. Transcriptional noise has a well-described role in promoting phenotypic diversification in homogeneous populations^24,25,26,27,28^ and we therefore reasoned that it could underlie the probabilistic awakening transitions observed in our lineage-tracing data.

Our analysis showed a sharp rise in transcriptional noise during awakening, peaking just before population regrowth (Figure 3). This enhances expression variability in cell identity genes, increasing the likelihood that dormant DTPs stochastically transition into proliferative phenotypes once the drug is removed. The elevated noise returned to pre-treatment levels once heterogeneity was restored, indicating that noise elevation is a transient property of the awakening window rather than a stable property of any clone or cell state. These dynamics define a narrow interval in which adaptation may be most vulnerable to intervention.

Clinical samples of patients that follow a similar pattern of response followed by relapse also display an enrichment of transcriptional noise following treatment and preceding relapse (Figure 4). These results provide direct experimental evidence in clinically relevant models that transcriptional noise drives plasticity-led adaptation, representing a mechanistic bridge between stochastic gene expression and evolutionary fitness. While previous studies proposed that noise facilitates stochastic fate transitions in stem or cancer cells, they have largely relied on mathematical models or limited experimental systems. For example, *Hang et al.* described that noise-mediated heterogeneity in Sca-I expression influences fate decisions in murine hematopoietic stem cells^72^ and *Gupta et al.* applied Markov modelling to show stochastic state transitions in breast cancer cells^73^. Our current study however directly links noise dynamics to tumour recovery and relapse, establishing transcriptional noise as a functional determinant of evolutionary potential.

Although we have not directly examined the contribution of specific genetic drivers (e.g. TERT, ATRX) to transcriptional noise, our models span different MYCN and ALK genotypes – SKNSH (MYCN^WT^, ALK^mut^), NB039 (MYCN^amp^, ALK^WT^), NB067 (MYCN^amp^, ALK^mut^) – yet display similar noise-driven plasticity dynamics. Moreover, we extend these observations to hepatoblastoma models (Figure 6; Figure S14–S15), a genetically distinct entity, suggesting that noise-mediated plasticity may represent a more generalized mechanism across paediatric cancers. Although the specific noisy genes differed between models, we identified conserved sets of transcriptional patterns: a shared core of noisy genes, which was consistently present across treatment stages, and an awakening-specific set that emerged post-treatment. These gene sets were both associated with the particular cell identity of the model: plasticity (i.e. EMT), cell migration, and neurogenesis in heterogeneous neuroblastoma cell lines; neurogenesis in adrenergic-enriched patient-derived models; and liver development and plasticity-related processes in hepatoblastoma. Together, these data support a model in which transcriptional noise affects lineage-determining gene networks to promote cell state transitions during both intrinsic and adaptive plasticity.

Our mechanistic analyses show that AP-1 sites become more accessible after treatment in preclinical models and patient samples, and that this accessibility persists at recovery (Figure 2C–D). Significantly, in a cohort of patients with longitudinal samples^47^, accessible chromatin around AP-1 motifs was associated with tumour-specific cell identities that tolerate induction therapy and are linked to worse outcomes, raising the possibility that AP-1-mediated chromatin states reflect elevated adaptive potential in clinical samples. We also show that the histone demethylase KDM5A is a key regulator of noise-dependent plasticity-led evolution: KDM5A binding increases at noisy, cell-identity genes after treatment (Figure 5B–F), leading to narrowing of H3K4me3 domains (Figure 5E), a chromatin signature associated with increased transcriptional variability. Notably, during awakening, KDM5A relocates to loci enriched for AP-1 binding sites (Figure 5D).

We functionally validated the role of AP-1 and KDM5 activity in plasticity-led recovery through genetic and pharmacological perturbation. Downregulation of ATF2, a transcription factor of the AP-1 family whose motif was enriched in both preclinical models and clinical samples after treatment, attenuated recovery after a strong cisplatin bottleneck compared to control cells (Figure 2E–F). Inhibition of AP-1 activity using T-5224 reduced the increase in transcriptional noise observed during awakening (Figure 6F, S10H and S13C). Similarly, inhibition of KDM5 with KDM5-C70 suppressed phenotypic plasticity, reduced transcriptional noise, and prevented the narrowing of H3K4me3 peaks, resulting in the blockade of tumour cell regrowth in both neuroblastoma and hepatoblastoma models (Figure 5 and 6D–E). Of note, KDM5 inhibition was effective in HepG2 cells, which did not display measurable noise dynamics under our experimental conditions, so noise-independent contributions of KDM5 activity to awakening cannot be excluded.

Together, these results support a model in which AP-1 primes KDM5A recruitment to cell identity genes and enhances transcriptional noise through chromatin rewiring. This model is supported with prior evidence that AP-1 mediates chromatin remodelling at cell-identity genes and that KDM5 family members preferentially bind AP-1-rich regions and interact with chromatin-remodelling machinery recruited by AP-1. More broadly, our results uncover a direct mechanistic link between chromatin remodelling and transcriptional noise regulation during plasticity-led cancer evolution. This aligns with prior work showing that histone methylation modulates gene-expression variability in embryonic stem cells, and with observations that KDM5 activity enhances transcriptomic heterogeneity and treatment resistance in ER⁺ breast cancer^52^. In this framework, AP-1-mediated chromatin accessibility defines regulatory landscapes that are subsequently exploited by KDM5 to modulate transcriptional noise, thereby facilitating phenotypic plasticity and adaptive evolution across cancer contexts.

Collectively, our findings demonstrate that paediatric cancer cells leverage plasticity to generate phenotypic diversity and enable awakening from DTP states after therapy through regulated mechanisms involving chromatin-dependent induction of transcriptional noise. This study provides, to our knowledge, the first experimental evidence directly linking transcriptional noise to phenotypic plasticity and relapse in cancer. Beyond advancing our mechanistic understanding of plasticity and adaptive evolution, these findings position noise as a novel non-genetic driver of plasticity, a key cancer hallmark. This has important implications for the identification of biomarkers associated with increased adaptive potential and malignant progression. Moreover, our results open new therapeutic avenues by targeting transcriptional noise and plasticity to prevent relapse and limit tumour evolution. Although the clinical application of KDM5 inhibitors has been constrained by toxicity concerns^77^, recent advances in targeted protein degradation technologies – such as proteolysis-targeting chimeras (PROTACs) – offer a promising strategy to overcome these limitations and translate our findings into clinical benefit^78–80^. Ultimately, targeting the regulatory mechanisms underpinning transcriptional noise may represent a novel approach to reducing cancer plasticity and improving long-term treatment outcomes.

## METHODS

### Cell lines and treatment

SK-N-SH human neuroblastoma cells were originally provided by L. Chesler, ICR, London. HuH6 human hepatoblastoma cells were provided by R. Kappler, LMU, Germany. HepG2 human hepatoblastoma cells were purchased from ECACC. All cell lines were cultured in DMEM (Thermo Fisher Scientific, USA) supplemented with 10% fetal bovine serum (FBS) (Thermo Fisher Scientific, USA), 100 IU penicillin and 100μg/mL streptomycin (Sigma-Aldrich, USA) at 37°C and 5% CO_2_. Unless specified otherwise, cells were treated with 50µM IdU (I7125, Sigma-Aldrich, USA), 100nM TGF-ß (240-B-010, Biotechne, USA), 30µM KDM5-C70 (HY-120400, MedChem Express, USA), 10µM T-5224 (HY-12270, MedChem Express, USA) or an IC50 concentration of cisplatin (CAY13119, Cambridge Bioscience, UK): 3.18µM for SK-N-SH, 11.1µM for HuH6 and 14.1µM HepG2. To achieve transient pausing of transcription, cells were treated with 200µM DRB (Sigma-Aldrich, USA), which was removed after 6 hours to promote normal transcription.

### Wound healing assay

SK-N-SH cells were plated in a tissue-culture 48 well-plate (Thermo Fisher Scientific, USA) at a density of 20,000 cells per cm^2^. After 24 hours, a scratch was done with a pipette tip and wells were washed twice with PBS to remove detached cells and provided with fresh media containing the desired drug. DMSO (D2650, Sigma-Aldrich, USA) was added at a final concentration of 0.25% v/v. IdU (I7125, Sigma-Aldrich, USA) was titrated at 10, 20 and 50µM; if not stated otherwise, the concentration used was 50µM. TGF-ß (240-B-010, Biotechne, USA) was supplied at a final concentration of 100nM. Cell migration was monitored using the IncuCyte S3 instrument (Sartorius, Germany) acquiring images every hour at 10X magnification. Measurements of wound area were done using an ImageJ plug-in^81^ and normalized to wound area of each condition at 0 hours.

### Flow cytometry and sorting

Treated cells were detached using TrypLE Express (Thermo Fisher Scientific, USA) and collected in 5ml polystyrene FACS tubes. Cells were washed in in 1XPBS + 1% Bovine Serum Albumin (Thermo Fisher Scientific, USA), here FB buffer, pelleted at 400xg for 5 minutes at 4°C and resuspended in 100μL FB. For experiments not including KDM5-C70, cells were stained with 2μL of anti-CD44 APC/Fire 750 antibody (103062, BioLegend, USA) for 30 min at 4°C. Cells were washed by centrifugation in FB and resuspended in 300μL FB containing 25ng/μL DAPI (130-111-570, Miltenyi, Germany). In experiments including KDM5-C70, washed cells were first stained with 1μL Zombie Red viability dye (423109, BioLegend, USA) for 10 min at room temperature and 2μL of anti-CD44 APC/Fire 750 antibody for 30 min at 4°C. Cells were then washed by centrifugation in FB and resuspended in 300μL FB. Cells were analyzed using a BD LSR II or Symphony A5 (BD Bioscience, USA) or sorted using a Symphony S6 (BD Bioscience, USA). Data was analyzed using the FlowJo software.

### Colony forming assay

For the colony forming assay cells were seeded at 1000 cells/well of a 6-well plate in complete media. Colony growth was monitored, and media was replenished once a week. Plates were imaged using the Incucyte S3 system (Sartorius), and images were analysed using the confluence tool, which generates the yellow segmentation mask visible in the output images. Segmentation parameters, including hole filling and area thresholds, were optimised per experiment and then held constant throughout the analysis. For colony number quantification, the Analyze Particles tool in Fiji (ImageJ2) was used.

### Immunofluorescence

For immunofluorescence staining, 2x 10^5 cells were seeded in a 24-well plate containing glass coverslips. The day after, cells were treated with Cisplatin for the indicated amount of time. After treatment, cells were fixed with 4% PFA for 30 minutes at 4°C. Cells were then permeabilized with 0.2% Triton X-100 for 10 minutes at RT. Cells were washed twice and incubated in PBS buffer containing 1% BSA and 2% FCS (IFF buffer) for 30 minutes at room temperature (RT). Cells were stained overnight at 4 °C with primary antibody diluted in IFF buffer (anti-CD44 IM7, 1:500, eBioscience; anti-gH2A.XpPhospho (Ser139) 2F3, 1:200, Biolegend). The following day, cells were washed three times with PBS and subjected to conjugated secondary antibody diluted in IFF buffer (anti-Mouse-AF488 A21202 Invitrogen, anti-Rat-AF647 A78947 Invitrogen) at 1:2000 for 2h at RT. Coverslips were mounted with DAPI Fluoromount-G mounting medium and imaged the with a confocal microscope (Zeiss LSM700).

### Barcoding of SK-N-SH cells

The CloneTracker library was a gift from Andrea Sottoriva (BCXP1M3RP-XSP, Cellecta). SK-N-SH cell lines were cultured in normal growth media and barcoded by lentiviral transduction according to manufacturer’s recommendations. Following lentiviral titration, 1 million neuroblastoma cells were transduced with a lentiviral library using a multiplicity of infection (MOI) of 0.1 (corresponding to 10% of cells transduced) to attain a single barcode per cell. 100,000 barcoded cells were FACS-sorted based on RFP-positivity and expanded. This population constitutes the “original population” or POT.

### Experimental design of single cell RNA sequencing of barcoded SK-N-SH cell line

Barcoded SK-N-SH cells were seeded equally into 2 treatment groups and divided across three replicates. The seeding density was adjusted based on each treatment and a total of 5M cells were seeded for each sample. Untreated cells were collected at confluency. The remaining three samples were exposed to cisplatin treatment (3.18 uM). After 1 week (T1), the treatment was removed and cells were detached, 10% of the total population was cryopreserved in four aliquots of 2.5% and the remaining cells were reseeded at a fixed density and grown under normal growth media for 1 week (T2) or 4 weeks (T3). Cell counts were determined via the Countess II Automatic Cell Counter (Thermo Fisher).

### Single cell RNA profiling of barcoded SK-N-SH cell line

After thawing, single cells were washed with PBS and were resuspended with a buffer (Ca++/Mg++ free PBS + 0.04% BSA). Cells were tagged with the CMO oligos (10X Genomics) as per manufacturer’s instructions. Samples were multiplexed in groups of 6, each containing 5000 cells/samples for a total of 30000 cells/pool.

Pooled single cell suspensions were loaded on a Chromium Single Cell 3′ Chip (10X Genomics) and were run in the Chromium Controller to generate single-cell gel bead-in-emulsions using the 10X genomics 3′ Chromium v2.0 platform as per manufacturer’s instructions. Single-cell RNA-seq and CMO libraries were prepared according to the manufacturer’s protocol and the library quality was confirmed with a TapeStation High-Sensitivity DNA Kit (Agilent, 5067-4627). Cellecta barcode libraries were prepared using 25% of the adaptor-ligated cDNA to amplify only the barcoded cDNA product. Barcodes were pre-amplified using the P5-Read1 primer (ACACTCTTTCCCTACACGACGCTCTTCCGATCT) and the P7-Adapter-FBPI Cellecta primer (GTGACTGGAGTTCAGACGTGTGCTCTTCCGATCTCCGACCACCGAACGCAACGCACGCA), following which Illumina indexes were added. Samples were sequenced on an Illumina NovaSeq 6000 according to standard 10X Genomics protocol, aiming for approximately 25,000 reads per cell for gene expression libraries and 1000 reads per cell for CMO/Cellecta barcode libraries.

### Single cell data pre-processing

10x Genomics CellRanger (v6.1.2) was used to process raw sequencing FASTQs, using cellranger count (or multi in the case of samples multiplexed with CellPlex) pipeline to conduct read alignment to human genome reference (GRCh38-2024-A), sample demultiplexing and generate feature-barcode matrix. For PDX models, read alignment was conducted using combined human and mouse genome reference (GRCh38_and_GRCm39-2024-A), to observe any mouse gene expression contamination within the sample.

EmptyDrops method from DropletUtils (v1.20.0) package^82,83^ was implemented on cellranger output H5 matrix to remove empty droplets with false discovery rate of 0.1%. Data files were merged and then normalised using logNormCounts from scuttle (v1.10.1) package^57^ for downstream QC and processing.

Cell-level filtering was carried out with the following thresholds: mitochondrial gene content <10%, ribosomal gene content >5%, and UMI counts >1000 for cell lines, mitochondrial gene content <40%, ribosomal gene content >0.3%, and UMI counts > 1000 for PDO models, mitochondrial gene content <30%, ribosomal gene content >0.2%, and UMI counts > 1000 for PDX models. Filtering at the gene-level was implemented to remove genes with fewer than five reads. Downstream normalisation and scaling were performed using Seurat (v4.3.0)^84^ with the SCTransform function, using the top 1000 variable genes, and regressing out cell cycle and mitochondrial genes. The SCTransform data output was used to perform a principal component analysis (PCA). The batch correction algorithm Harmony (v0.1.1) was applied across CMO multiplexed pools (where appropriate), and the output used for Uniform Manifold Approximation and Projection (UMAP)-based dimensional reduction. Clustering was carried out using the Seurat FindClusters function and gene expression markers for the clusters were identified by a Wilcoxon Rank Sum test, using the Seurat FindAllMarkers function. Gene set enrichment analysis using clusterProfiler (v4.6.2)^85,86^ was then carried out on the cluster markers. To compare changes in cell clusters, statistical testing was conducted using student t-test between each sample and T0 (untreated), with adjusted p-value calculated using Bonferroni method for multiple-comparison correction.

### Cell state score generation

Single-cell expression signatures for the ADRN and MES cell states were created using the AddModuleScore function in Seurat and published gene sets^40^. To calculate the cell state score we used a method based on that used by Tan et al.^87^. Statistical testing (t-tests) was used to determine whether differences between ADRN and MES gene set expression were significant. The t-statistic was taken as the cell state score and the statistical significance was further used to segregate cells into phenotypic groups: ADRN (AMT score < 0; P < 0.05), intermediate ADRN (AMT score < 0; P ≥ 0.05), intermediate MES (AMT score > 0; P ≥ 0.05) and MES (AMT score > 0; P < 0.05). The two intermediate classifications were grouped together as INT. To compare changes in phenotypic cell state assignment, statistical testing was conducted using student t-test with adjusted p-value calculated using Bonferroni method for multiple-comparison correction.

### Cellecta barcodes amplification and next generation library preparation

SK-N-SH cells were harvested and pelleted at the relevant timepoints. Genomic DNA was isolated using AllPrep kit (Qiagen) according to manufacturer’s instructions and quantified using Qubit (Life Technologies). Amplicon PCR was conducted using 50ng of DNA (or 20ng for samples with lower DNA yield) along with 2x Accuzyme mix (Bioline) to amplify barcodes using the following primer sequences (Forward - ACTGACTGCAGTCTGAGTCTGACAG, Reverse - CTAGCATAGAGTGCGTAGCTCTGCT). After identifying and purifying PCR product (consisting of 80-bp product + 30-bp semi-random barcode), NGS libraries were prepared using NEBnext Ultra II DNA library preparation kit for Illumina (New England Biolabs) and quantified with using a TapeStation (Agilent Genomics). Samples were balanced and pooled together to have final molarity of 4nM, NGS was then performed by ICR Genomics facility (TPU) using NovaSeq SP PE150.

### Cellecta barcodes bioinformatics analysis

FASTQ files from bulk experiments were mapped to a custom index reference generated from white-list (all possible sequences) of Cellecta barcodes, using Burrows-Wheeler aligner (BWA-MEM tool; version 0.7.17)^88^. Once mapped BAM files were generated using Sequence Alignmnet/Map (SamTools; v1.18)^89^. Gene counts were quantified using FeatureCounts from Rsubread. Parameters were set to filter out reads with mapping quality less than 30 (minMQS = 30).

To observe lineage dynamics, representative percentage was calculated for each lineage within each sample (frequency of Cellecta barcode / frequency of all Cellecta barcodes x 100).

Shannon diversity index^90^ of all DNA barcodes from each experimental samples was calculated between conditions to observe changes in lineage diversity as per following equation: H=−∑[(pi)×log(pi)], where pi is proportion of individual lineage within the population. Statistical significance of results was determined using pairwise *t-tests*.

Cumulative proportion quantification of all DNA barcodes from each sample was calculated by ranking all barcodes from most to least abundant (sum of frequency). Cumulative proportion was then calculated as cumulative sum of barcode abundance / sum of abundance. For statistical quantification an area under the curve (AUC) value was calculated for each sample using *trapz()* (from R package prcama) of cumulative proportion and abundance from cumulative proportion plot. Statistical significance of results was determined using pairwise *t-tests* between untreated and each other samples.

For top 100 POT barcode analysis, barcodes in POT sample were ranked most to least frequent and the top 100 most abundant barcodes were extracted. Remaining experimental samples were filtered for these barcodes and visualised.

For top 100 recovery barcode analysis, barcodes in each T3 sample were ranked most to least frequent and the top 100 most abundant barcodes were extracted. Samples from the same experimental treatment arm were filtered for these barcodes and visualised. Common colour scheme for barcodes of interest at T3 was generated across all top 100 barcodes lists from all experimental arms, to allow visualisation of common barcodes across experimental arms.

For scRNA-seq experiments Cellecta barcodes from 10x Chromium libraries were isolated using STAR solo^91^ with the same white list of barcodes used for BWA for bulk experiments. Where multiple barcodes were observed within a single cell, cells were assigned the barcode with the higher proportion of representation from read counts. To account for sparse data, lineages were filtered to ensure each barcodes were observed at least 3 times within the entire dataset.

Plasticity transition dynamics analysis was conducted by assigning each cell within each lineage a plasticity score based on the number of cell states occupied; either =1 for a single phenotypic state or > 1 for multiple phenotypic states. For analysis frequency of cells in each group were aggregated across replicates. Chi-squared statistical testing was conducted in a pairwise fashion to observe whether there was an association between sample and plasticity based on frequency of each plasticity group. Statistical test was conducted between untreated and each other sample and subject to multiple comparison correction using Bonferroni’s method. Directionality of statistical significance was calculated as (frequency of cells in >1 plasticity group / total cell frequency in sample) - (frequency of cells in >1 plasticity group from untreated / total cell frequency in untreated). Positive was termed change value > 0 and negative < 0.

To identify and track cellular lineages that successfully persisted throughout the treatment, we implemented a matched clone analysis focusing on clones present in both untreated and recovery conditions. We identified winner clones by filtering criteria. Only clones with a minimum of 3 cells in both the untreated condition and the 4-week off-treatment timepoint were included in the analysis. For each matched clone, we tracked their AMT state transitions across all time points. The dominant cellular state at each timepoint was determined by the most frequent annotation among cells within each clone. To maintain consistency and statistical reliability, the minimum cell threshold of 3 cells per condition was applied across all timepoints. Clone trajectories were visualized using alluvial flow diagrams implemented with the ggalluvial R package, enabling tracking of individual clone over time.

Lineage diversity was quantified as within-replicate (α) and between-replicate (β = γ/α) diversity using Hill numbers of order q = 1, adapted from Whiting et al.^46^.

### Patient-derived xenograft establishment

PDX (IC-pPDX-112 and GR-NB5) were established from neuroblastoma tumor biopsies obtained at diagnosis, as described previously, following informed consent of parents or guardians^92,93^. This study was performed in accordance with the recommendations of the European Community (2010/63/UE) for the care and use of laboratory animals. Experimental procedures were approved by the ethics committee of the Institut Curie CEEA-IC #118 (Authorization APAFIS#11206-2017090816044613-v2) in compliance with the international guidelines. The establishment of PDX received approval by the Institut Curie institutional review board OBS170323 CPP ref 3272; n° de dossier 2015-A00464-45. In brief, tumor samples were engrafted subcutaneously in immunocompromised mice. Upon successful establishment (> P2) and upon reaching ethical size, PDX tumors were cryopreserved for banking, morphological and molecular analyses according to procedures described previously.

### Spheroid morphology quantification

IC-pPDX-112 was dissociated as previously described (*63*) and cultured in DMEM supplemented with 10% FBS. 80,000 cells/well were seeded in a tissue-culture flat-bottom 96-well plate (Thermo Fisher Scientific, USA). After 10 days in culture, cells were treated with DMSO (0.25% v/v) or IdU (50µM) in the presence or absence of TGF-ß (100nM) and monitored using the IncuCyte ZOOM instrument (Sartorius, Germany) at a 10X magnification every hour. To analyze their morphology, spheroids were manually segmented, and area, roundness and total number of fusion events were calculated using ImageJ.

### RNA sequencing of PDX

IC-pPDX-112 was dissociated as previously described^94^ and cultured in DMEM supplemented with 10% FBS. 1×106 cells/well were seeded in tissue-culture 6-well plates. After 10 days in culture, cells were treated with DMSO (0.25% v/v) or IdU (50µM) in the presence or absence of TGF-ß (100nM). 5 days after treatment cells were collected and RNA was isolated using the RNeasy Micro Kit (Qiagen, Germany). RNA quality was assessed using the Bioanalyzer System (Agilent, USA) and 600ng of RNA were used for library preparation with the Illumina Stranded mRNA Prep, Ligation kit (Illumina, USA).

Sequencing was performed in a NovaSeq instrument (Illumina, USA) using 30 million clusters per sample and PE100 reads. Pre-processing of the raw sequencing reads was performed using an in-house pipeline available via https://zenodo.org/records/13744441. Briefly, reads were trimmed using TrimGalore v0.6.2 to remove adapters and quality-checked using FastQC v.0.11.9. Reads aligning to the mm10 murine reference genome were removed using xengsort (*64*) to avoid contamination from host mouse cells. Remaining reads were aligned to the GRCh37/hg19 human reference genome using STAR aligner v2.7.6 and gene-level counts were quantified with featureCounts using GENCODE annotation. Differential gene expression analysis was performed using DESeq2 (v1.40.2) in R (v4.3.0).

### Patient-derived models and treatment for scRNA-seq analysis

The PDX line GR-NB5 (MYCN^AMP^, previously named MAPGR-B25-NB-117^41^) was originally provided by G. Schleiermacher (Curie Institute, Paris, France). Cells were obtained from enzymatical dissociation of the PDX and cells were cultures in in low-Glucose GlutaMAX DMEM (Thermo Fisher) supplemented with 25% Ham’s F-12 Nutrient Mix, 2% FBS (Thermo Fisher), B-27 Supplement (50X) (Thermo Fisher), minus vitamin A N-2 Supplement (100X) (Thermo Fisher), 100 U/mL penicillin-streptomycin (Sigma), 20 ng/mL Animal-Free Recombinant Human EGF (Peprotech), 40 ng/mL Recombinant Human FGF-basic (Peprotech), 200 ng/mL Recombinant Human IGF-I (Peprotech), 10 ng/mL Recombinant Human PDGF-AA (Peprotech) and 10 ng/mL Recombinant Human PDGF-BB (Peprotech). NB039 (MYCN^AMP^ ALK^WT^) and NB067 (MYCN^AMP^ ALK^R1275Q^) patient-derived spheroids were originally provided by J. Molenaar (Princess Maxima Centrum, Netherlands). PDOs were cultured in low-Glucose GlutaMAX DMEM (Thermo Fisher) supplemented with 25% Ham’s F-12 Nutrient Mix, B-27 Supplement (50X) (Thermo Fisher), minus vitamin A N-2 Supplement (100X) (Thermo Fisher), 100 U/mL penicillin-streptomycin (Sigma), 20 ng/mL Animal-Free Recombinant Human EGF (Peprotech), 40 ng/mL Recombinant Human FGF-basic (Peprotech), 200 ng/mL Recombinant Human IGF-I (Peprotech), 10 ng/mL Recombinant Human PDGF-AA (Peprotech) and 10 ng/mL Recombinant Human PDGF-BB (Peprotech). After thawing, single cells were washed with PBS and were loaded onto a Chromium Single Cell 3′ Chip (10X Genomics) and were run in the Chromium Controller to generate single-cell gel bead-in-emulsions using the 10X genomics 3′ Chromium v2.0 platform as per manufacturer’s instructions. Single-cell RNA-seq libraries were prepared according to the manufacturer’s protocol and the library quality was confirmed with a TapeStation High-Sensitivity DNA Kit (Agilent, 5067-5585). Samples were sequenced on an Illumina NovaSeq 6000 according to standard 10X Genomics protocol, aiming for approximately 50,000 reads per cell for gene expression libraries and 1000 reads per cell for CMO/Cellecta barcode libraries.

### snRNA-seq of clinical samples

Six matched samples at diagnosis and post-treatment were obtained form VIVO biobank (Newcastle, UK; Table S5), supported by Cancer Research UK & Blood Cancer UK (Grant no. CRCPSC-Dec21\100003).

Nuclei were isolated from fresh frozen samples by mincing in a Tween-based lysis buffer containing RNase inhibitors, washed, counted on a LUNA fluorescence cell counter (Logos Biosystems) and prepared for the 10X Chromium using the Chromium Next GEM Single Cell 3ʹ Kit v4, targeting 20,000 nuclei per sample. cDNA and libraries were prepared for sequencing with the same Chromium kit and sequenced (PE150) on a NovaSeq X (Illumina). Three matched samples and two pre-treatment samples were successfully included in the snRNA-seq run (Table S5).

Raw sequencing reads were aligned to the human reference genome (GRCh38) using Cell Ranger. True cells were called using EmptyDrops (0.1% FDR; ≥200 genes; ≥500 UMIs), and doublets removed with scDblFinder (4% expected rate). Cells were further filtered to retain those with ≥200 detected genes, <10% mitochondrial content, and >0.1% ribosomal content. Gene expression was normalised using SCTransform, regressing out cell cycle score and mitochondrial percentage. Dimensionality reduction was performed by PCA (30 components) followed by Harmony batch correction across patients, UMAP visualisation, and unsupervised clustering at resolution 0.2. Cells were annotated to cell types by expression of previously identified markers or neuroblastoma tumours and stroma^95^.

Malignant cells were identified by CNV inference using InferCNV (clusters 11 and 16 as normal reference; cutoff 0.1). Malignant neuroendocrine clusters were collapsed and further annotated based on similarity of expression of makers of sympatho-adrenal development^39^.

### MuTrans analysis

Pre-processing of the scRNA-seq data, split by drug condition and experimental timepoint, was carried out analogous to as outlined above using Scanpy (v1.9.3)^96^. Using the filtered data as inputs, the count matrices were normalised and log transformed with a pseudo-count of 1. The number of transcripts per cell and the mitochondrial content were regressed out and the data was scaled, prior to PCA. Dimensionality reduction using UMAP and Leiden clustering was carried out.

A neighbourhood graph was computed for the processed data using scanpy.pp.neighbors (metric = “cosine”). Construction of cell-fate dynamical manifolds and downstream analysis of cell-state transition was then performed using the MUltiscale method for TRANSient cells^45^. Attractor wells were annotated into “plastic” or “committed” based on median entropy across all cells in the well. Phenotypic annotation into ADRN or MES represents the most common cell identity in a well. Wells that contained the highest percentage of BRI cells were assigned this phenotype.

### Transcriptional noise quantification

Single-cell RNA-seq data on HuH6 and HepG2 was obtained from https://dataview.ncbi.nlm.nih.gov/object/PRJNA1247955?reviewer=g4kgeej9of8fb9sf5ng8hdepnl and pre-processed as previously described. Processed expression matrices were then analyzed in R studio (v4.3.0) using the BASiCS package (v2.12.3)^49,50^. For analysis of plasticity during cisplatin recovery, mesenchymal/reservoir cells were split into equally sized groups based on timepoint. Group size was determined by the smallest sample. Markov chain Monte Carlo simulations were performed using BASiCS_MCMC function to estimate expression parameters (N = 4000, Thin = 10, Burn = 2000). Chain convergence was visually inspected. For differential noise analysis between conditions BASiCS_TestDE function was used with default parameters. Identification of highly variable genes (HVG) within specific conditions was performed using BASiCS_DetectHVG function with a VarThreshold of 0.6. For the snSK-N-SH of SK-N-SH of cells treated with DMSO, KDM5-C70 or T-5225 VarThreshold was reduced to 0.3 due to low HVG recovery.

### Transcriptional bursting inference

Transcriptional burst kinetics (burst size (s), burst frequency (f), and duty cycle (d)) were inferred for MES cells at the Persistence versus Awakening timepoints using D3E^51^, which fits a two-state promoter (Poisson–Beta) model to single-cell expression count distributions. Analysis was performed separately for each of three biological replicates, and per-gene parameter estimates and associated p-values (pSize, pFreq, pDuty) were averaged across replicates prior to downstream analysis. Multiple testing correction was applied using the Benjamini–Hochberg procedure independently within each replicate before averaging. Genes for which the goodness-of-fit statistic (GOF) indicated model failure in either condition were excluded. To characterise global shifts in bursting kinetics between timepoints, absolute log2 fold-changes in s, f and d were compared across tertiles of delta HVG probability (Awakening - Persistence) using Wilcoxon rank-sum tests with pairwise comparisons between bins.

Each gene was assigned to one of eight burst-pattern categories based on the direction of change (up/down) of each parameter (s, f, d) between Persistence and Awakening. Enrichment of high-variability gene status (defined by BASiCS posterior HVG probability in Awakening) within each pattern was assessed by Fisher’s exact test, with odds ratios and Benjamini– Hochberg-adjusted p-values reported. Schematic mRNA traces for selected patterns were generated from a deterministic two-state promoter simulation (dM/dt = k(t) − γM(t), with constant degradation rate γ and production rate k active during the ON state), parameterised using the mean fold-changes in s, f and d observed for each pattern relative to baseline persistence parameters set to unity. These traces are illustrative of the qualitative bursting dynamics implied by each pattern and do not constitute fits to the observed single-cell data.

### Generation of d_2_GFP cell lines

To construct the destabilized GFP (d_2_GFP) reporter, the d_2_GFP gene was amplified by PCR from the Lentiviral-TOP-dGFP-reporter plasmid (Addgene plasmid #14715) and cloned downstream of an EF1a promoter in the lentiviral plasmid pLX307 Luciferase (Addgene plasmid #117734) using the NEBuilder HiFi DNA Assembly Cloning kit (E5520S, New England Biolabs, USA). Lentiviral-TOP-dGFP-reporter was a gift from Tannishtha Reya and pLX307 Luciferase was a gift from William Hahn. Lentiviral particles were produced in HEK293T cells using psPAX2 and pMD2.G as packaging vectors and diluted 1:1 with fresh DMEM to infect SK-N-SH cells. Transduced cells were selected with 1µg/mL puromycin (P33020, Melford Labs, UK) until complete eradication of control untransduced cells was achieved. The remaining d_2_GFP-positive cells were then sorted using a Symphony S6 instrument (BD Biosciences, USA) to increase purity.

### Drug screen

SK-N-SH d_2_GFP were seeded in duplicated in a 6-well plate at a density of 10,000 cells per cm^2^. After 24h, cells were treated with IdU cisplatin (50μM), KDM5-C70 (30μM) or cisplatin (3.18μM). Five days after treatment, the expression of CD44 and d_2_GFP was assessed by flow cytometry. For analysis, 2000 CD44^high^ cells were sampled and d_2_GFP expression in each cell was normalized to the average expression across the sample.

### KDM5-C70 treatment during recovery from cisplatin

For SK-N-SH, control untreated cells were plated at a seeding density of 10,000 cells per cm^2^, while treated cells were seeded at 20,000 cells per cm^2^ in a 6-well plate in triplicate. After 24h, cells were treated with cisplatin (3.18μM) for seven days, after which cisplatin was washed and KDM5-C70 (30μM) was added only once or weekly for a month. The phenotypic composition of the population was assessed by CD44 flow cytometry at 0, 7, 14 and 35 days post cisplatin treatment.

For live imaging and growth assessment in hepatoblastoma models, 50,000 cells were seeded in duplicate or triplicates in a 24-well plate. After 24h, cells were treated with an IC50 dose of cisplatin for 48h. After that period, cisplatin was washed, KDM5-C70 (30μM) or DMSO was added once and cells were followed until normal growth was achieved. Cell growth was monitored and quantified using the IncuCyte S3 system (Sartorius, Germany).

For the colony-forming assay 200,000 HepG2 cells were seeded in duplicates in a 6-well plate. After 24h, cells were treated with cisplatin (14.1μM) for seven days. After that period, cisplatin was washed, KDM5-C70 (30μM) or DMSO was added once and cells were kept in culture with weekly media changes until day 35 post cisplatin treatment. On day 35, cells were imaged with the IncuCyte S3 system to quantify colony diameter. After this, media was aspirated, cells were washed with 1xPBS, stained with GIEMSA dye (Sigma-Aldrich, USA) and imaged to quantify number of colonies.

### Western blotting

1×10^6^ cells were harvested and lysed using RIPA buffer (Thermo Fisher Scientific, USA) supplemented with proteinase and phosphatase inhibitors. Lysates were centrifuged at 13,000xg for 20 min to isolate the proteome. Protein lysates were quantified using the DC Protein Assay kit (Bio-Rad, USA). 30µg of protein were denaturalized by incubating at 95°C for 5min in the presence of 2-mercaptoethanol. Samples were loaded in a Bolt 4-12% Bis-Tris Plus Gel (Thermo Fisher Scientific, USA), run using MOPS buffer and transferred into a PVDP membrane. Membranes were blotted using anti-KDM5A (Jarid1A/RBBP2 (EPR18651) Rabbit mAb, ab194286, Abcam), anti-KDM5B (JARID1B (E2X6N) Rabbit mAb, 15327, Cell Signalling Technology) or anti-GAPDH (GAPDH (14C10) Rabbit mAb, 2118, Cell Signalling Technology) primary antibodies and anti-Rabbit IgG-HRP (Goat Anti-Rabbit Immunoglobulins/HRP, P044801-2, Agilent Technologies, UK) secondary antibody at the recommended concentrations.

### Dose response curve

Cell lines were seeded at 2000 cells/well in 384-well plates and cultured for 24 hours prior to dosing. Cells were dosed using a 7-dose response curved with three technical replicates per concentration. Cell viability was assessed using CellTiterGlo-3D (Promega, USA) at day 3 after dosing. Cell viability was normalized to background and solvent controls, and dose response curves were fitted using isotonic regression, as previously described^97^.

### CUT&Tag sample preparation and analysis

SK-N-SH cells were seeded in duplicate in a 6-well plate and treated with an IC50 concentration of cisplatin or left untreated. After one week, untreated and treated cells were collected and viably frozen (“Untreated” and “Persistence” timepoints). For the remaining cells cisplatin treatment was removed. One arm was left to recover untreated and a second arm was treated with 30μM KDM5-C70. Cells were collected and frozen a week later (“Awakening” and “Awakening + KDM5-C70”, respectively). Samples were processed using the Cut & Tag-IT Assay Kit (Active Motif, USA) as per manufacturer’s instructions with 1μg of antibody: anti-KDM5A (Jarid1A/RBBP2 (EPR18651) Rabbit mAb, ab194286, Abcam), anti-KDM5B (JARID1B (E2X6N) Rabbit mAb, 15327, Cell Signalling Technology), anti-H3K4me3 (Tri-Methyl-Histone H3 (Lys4) (C42D8) Rabbit mAb, 9751S, Cell Signalling Technology). Sequencing was performed in a NovaX instrument using 50 paired-end cycles.

For H3K27ac CUT&Tag, cells were only collected at the “Untreated” stage and processed using the same protocol with an anti-H3K27ac antibody (Rabbit pAb, C15410196, Diagenode). CUT&Tag sequencing data were processed following the computational pipeline described by *Zheng et al.*^98^, with modifications for time series analysis. Raw paired-end sequencing reads were aligned to the human reference genome (GRCh38) using Bowtie2 (v2.4.2) with CUT&Tag-optimized parameters (--local --very-sensitive --no-unal --no-mixed --no-discordant-I 10-X 700). Aligned reads were coordinate-sorted and indexed using SAMtools (v1.11). Quality control metrics including alignment rates, duplication rates, and library complexity were assessed for all samples using custom scripts adapted from the tutorial.

Duplicate reads were removed using Picard MarkDuplicates (v2.23.8) with REMOVE_DUPLICATES=true. Peak calling was performed using MACS2 (v2.2.7.1) in paired-end mode (-f BAMPE) with a q-value threshold of 0.1, using untreated samples as matched controls for each histone mark. Peaks from the two replicates of each condition were combined as the union of the replicate peak sets. For heatmaps, peaks from all treated conditions and replicates were merged with HOMER mergePeaks (-d 7500).

For quantitative visualization, normalized coverage tracks were generated using bamCoverage with RPGC normalization (effective genome size: 2,913,022,398 bp) and 10 bp binning. Signal profiles and heatmaps were created using deepTools computeMatrix and plotHeatmap, with analyses centred on transcription start sites (TSS) and peak summits using ±3 kb flanking regions. Gene-specific chromatin dynamics were assessed by assigning peaks to the nearest gene within 5 kb using bedtools window to evaluate lineage-specific chromatin remodeling across treatment time points.

Genomic annotation of CUT&Tag peaks was performed using HOMER (v4.11) annotatePeaks.pl against GRCh38. Peaks were classified into five regulatory classes combining TSS distance, chromatin co-occupancy (bedtools intersect), and external enhancer annotation: Proximal Active (<3 kb, H3K4me3+ or H3K27ac+), Proximal Other (<3 kb, no marks), Distal Active Enhancer (≥3 kb, H3K27ac+), Distal Putative (≥3 kb, EnhancerAtlas 2.0 SK-N-SH+, H3K27ac−), and Distal Other (≥3 kb, no marks). H3K27ac peaks were identified from untreated SK-N-SH CUT&Tag. De novo and known motif enrichment on each class was performed using HOMER findMotifsGenome.pl (hg38, 200 bp window, repeat-masked).

### CUT&RUN processing and peak calling

SK-N-SH cells were seeded in duplicate in a 6-well plate and treated with an IC50 concentration of cisplatin. After one week, cisplatin treatment was removed. One arm was left to recover untreated and two other arms were treated with 30μM KDM5-C70 or 10μM T-5224. Cells were collected and frozen a week later (“Awakening”, “Awakening + KDM5-C70”, “Awakening + T-5224”).

Samples were processed using the ChIC/CUT&RUN Assay Kit (Active Motif, USA) as per manufacturer’s instructions with anti-KDM5A (Jarid1A/RBBP2 (EPR18651) Rabbit mAb, ab194286, Abcam), anti-H3K4me3 (Active Motif), anti-ATF2 (E243, ab32160, Abcam), anti-cJun (60A8, 9165S, Cell Signalling Technologies), anti-cFos (9F6, 2250S, Cell Signalling Technologies). Libraries were prepared using Active Motif’s DNA Library Kit For Illumina. Sequencing was performed in a NovaSeq X instrument using 38 paired-end reads, targeting 5M read pairs per sample. Adapters were trimmed using Cutadapt v4.9 (TruSeq adapters; -- nextseq-trim=20-O 1-m 20;)^99^, and reads were aligned to GRCh38 using Bowtie2 with the CUT&Tag parameters. Duplicates were removed using Picard v2.23.8, and the two replicates per antibody and condition were pooled using SAMtools v1.11 to improve binding-site recovery^100^. Peaks were called using MACS3 v3.0.4 (-f BAMPE-g hs-q 0.1 --nomodel --keep-dup all), without a control and with local background estimation. Peaks overlapping the ENCODE hg38 blacklist v2 were excluded.

CPM-normalised coverage tracks were generated from pooled deduplicated BAMs using deepTools bamCoverage v3.5.5 (--binSize 50 --extendReads), excluding blacklist regions. CPM normalisation was selected based on CUT&RUN benchmarking^101^. Mean CPM for H3K4me3, KDM5A, ATF2, Fos and Jun was quantified with multiBigwigSummary for each of 40,670 H3K4me3-marked promoter intervals from the SK-N-SH CUT&Tag map. Signals used for ranking and matching were averaged across Awakening conditions.

To assess KDM5A enrichment at AP-1-high promoters, the top H3K4me3 quartile (10,168 promoters) was retained, and the top and bottom quartiles for each AP-1 factor within this set defined AP-1-high and AP-1-low groups (2,542 promoters each). KDM5A and H3K4me3 were compared in each condition as ratios of median CPM, with the ratio of median H3K4me3 used as a balance check.

### Single nuclei multiomics

We performed single-nuclei multiome sequencing (snRNA-seq and snATAC-seq) on SK-N-SH cells pre-and post-cisplatin treatment (one untreated and one recovered sample). Viable samples were prepared for sequencing following the 10x Genomics Multiome (snRNA-seq and snATAC-seq) protocol.

10x Genomics CellRanger-Arc (v2.0.2) was used to process raw sequencing FASTQs, using cellranger-arc count pipeline to conduct read alignment to human genome reference (GRCh38-2024-A) for both snRNA-seq and snATAC-seq to generate feature-barcode matrices and ATAC-seq fragment files.

### Single nuclei RNA preprocessing and visualization

EmptyDrops method from DropletUtils (v1.20.0) package^82,83^ was implemented on H5 matrix to remove empty droplets with false discovery rate of 0.1%. Data files were merged and then normalised using logNormCounts from scuttle (v1.10.1) package^102^ for downstream QC and processing. Feature-barcode gene expression matrices were filtered to remove low quality nuclei (> 50 UMI detected counts, > 0.25% ribosomal reads and < 40% mitochondrial gene content).

Filtering at the gene-level was implemented to remove genes with fewer than three reads. Downstream normalisation and scaling were performed using Seurat (v4.3.0)^84^ with the SCTransform function, using the top 2000 variable genes, and regressing out cell cycle and mitochondrial genes. The SCTransform data output was used to perform a principal component analysis (PCA). Uniform Manifold Approximation and Projection (UMAP)-based dimensional reduction. Clustering was carried out using the Seurat FindClusters function and gene expression markers for the clusters were identified by a Wilcoxon Rank Sum test, using the Seurat FindAllMarkers function. Gene set enrichment analysis using clusterProfiler (v4.6.2)^85,86^ was then carried out on the cluster markers. To compare changes in cell clusters, statistical testing was conducted using student t-test between either ADRN cells pre-and post-treatment, or between plastic and non-plastic MES cells, with adjusted p-value calculated using Bonferroni method for multiple-comparison correction. Cell state score generation was conducted as described previously.

### Single nuclei ATAC preprocessing and visualization

ArchR (v1.0.3)^103^ package was used to analyse snATAC-seq data. The distribution of the number of fragments per nuclei and transcription start site (TSS) enrichment was filtered ad hoc in each sample to retain data which was most likely representative of viable cells (> 1000 fragments and > 4 TSS). Following QC, using ArchR arrow files were generated with GrCh38/hg38 genome reference and filtered for doublets using filterRatio of 1.2. To visualise proximity of single nucleus chromatin accessibility profiles, we applied addIterativeLSI function to perform TF-IDF normalization, results were then projected onto UMAP (using 30 nearest neighbours).

### Single nuclei ATAC peak calling

Signac v1.6 CallPeaks function^104^ (which implements MACS3 peak caller^105^) was used to identify chromatin accessibility peaks in each sample. Blacklisted peaks and sex chromosome peaks were removed, resulting peaks were then combined with reduce function (GenomicRanges package^106^) which merges intersecting peaks.

### Single nuclei chromatin accessibility and transcription factor binding motif analysis

Combined dataset was filtered for cells which had both snATAC-seq and sn-RNA-seq following QC and peak calling. Using ArchR MarkerFeatures we identified differentially accessible peaks between samples of the same phenotypic state defined by gene expression profiles. Differentially accessible peaks were projected onto volcano plots. Transcription factor motif enrichment in differential peaks was calculated using cisbp database (http://cisbp.ccbr.utoronto.ca/^107^) version 2 to define motifs.

### siRNA transfection

SK-N-SH cells were seeded into six-well plates and transfected with ON-TARGETplus Human ATF2 SMARTpool or non-targeting pool. The transfection reagent used was the DharmaFECT 1 Transfection Reagent (Horizon Discovery Biosciences). Media containing the transfection reagents was changed after 6 hours and cells were analysed by immunoblotting and Flow Cytometry.

### Annotation of hepatoblastoma cells

The annotation of HuH6 and HepG2 hepatoblastoma cells was transferred from the ones defined by Chen et al^57^. For HuH6 clusters 1 and 2 were renamed PROL due to their differentiated and proliferative nature and their sensitivity to treatment; cluster 5 was labelled BRI for their intermediate position in the UMAPs, their progenitor-like markers and their dominance during awakening; and cluster 4 was renamed RES due to their progenitor nature and resistance to treatment. Newly generated snRNA-seq data on HuH6 was annotated as previously described^57^, using markers of liver development.

### Data availability

The raw single-cell RNA sequencing data used in this study can be accessed through the National Centre for Biotechnology Information - Sequence Read Archive (NCBI - SRA) using the following links: https://dataview.ncbi.nlm.nih.gov/object/PRJNA1256291?reviewer=ockms98q7ckoirf9hj0vmgtnn6 https://dataview.ncbi.nlm.nih.gov/object/PRJNA1247955?reviewer=g4kgeej9of8fb9sf5ng8hdepnl.

Previously published neuroblastoma plasticity datasets are available under BioProject PRJNA1071370.

The raw bulk RNA sequencing data on the PDX cells is available under restricted access, through the European Genome-Phenome Archive (EGA) with ID EGAD50000001770.

Processed data supporting this study objects have been deposited in Zenodo under accession code https://doi.org/10.5281/zenodo.23081009. Files are under restricted access until publication and are available to reviewers via the following link:

https://zenodo.org/records/23081009?token=eyJhbGciOiJIUzUxMiJ9.eyJpZCI6ImRiYWUzMzY2LTdjYzQtNGE4MS05MDVjLTVkOWVkNGYwYjZjNyIsImRhdGEiOnt9LCJyYW5kb20iOiJkMWE5ZThmNGY1OWFmZDQxYmE2NzFlMjljYjdkODRjMiJ9.hjyza8DW4bgODw16ccjbKd2oYsQwQmgaW5zyfLtalJC3ZKRqmXmv3mgljMCnoMRhUkCd4ERc9l39pjep1wrqIQ

## Code availability

Code used to process the data in this study is available in the following GitHub repository: https://github.com/PCEBrunaLab/AlloAnido-Roux-et-al-2026.

## ACKNOWLEDGEMENTS

We thank Katerina Rekopoulou and Nik Matthews for assistance with sample and library preparation for single-cell RNA-sequencing; Jan Molenaar for providing the patient-derived spheroid lines; Andrea Sottoriva for sharing the Cellecta barcode library and expertise in barcoding the cells as well as Salvatore Milite and Haider Tari for assistance and technical advice in analysis and barcode sequencing processing; Yura Grabovska and Chris Jones for technical expertise in the analysis of scRNA-seq data; Leor Weinberger, Melanie Beckett, Elizabeth R. Tucker, Evon Poon and Karen Barker for helpful discussion and technical advice; Peijie Zhou for help with the MuTrans analysis; Oscar M. Rueda and Maurizio Callari for technical advice regarding WGS and WES analyses. The PDX models were developed within the MAPPYACTS study following informed consent from patients and legal representatives. The authors thank all patients and parents/caregivers who participated in the Mappyacts trial and consented to the development of preclinical models. We acknowledge the ICR facilities: Light Microscopy (ICR-Services/Light-Microscopy-Facility: v1.8), Flow Cytometry (ICR-Services/ICR-Flow-Cytometry-Facility: v1.5), Genomics (ICR-Services/Genomics-Facility: v2.3) and Scientific Computing. We acknowledge the RSE Group at The Institute of Cancer Research for providing software enhancements, specifically Rachel Alcraft. Their support was instrumental in achieving the results presented in this paper. A.B. is supported by ICR London. This study was supported by Little Princess Trust Innovation Grant LPTINS2\9 (to A.B. and L.C.) and VAGABOND Marie Sklodowska-Curie Actions ITN (Horizon 2020) Grant 956285 (A.B., L.C. and G.S.). For the purpose of Open Access, the authors have applied a CC BY public copyright licence to any Author Accepted Manuscript (AAM) version arising from this submission.

## AUTHOR CONTRIBUTIONS

Methodology, A.A.A., C.R., S.H., A.S., E.C., C.E., A.S.S., C.B., A.Be.; experimental design, A.A.A., C.R., S.H., A.S., M.D.M., A.B..; analytical investigation, A.A.A., S.H., E.C., A.S.S., H.C., A.B.; writing–original draft, A.A.A., C.R., A.B.; writing–review and editing, A.A.A., C.R., S.H., A.S., E.C., C.E., M.D.M., A.B.; resources, L.C., G.S., B.G.; conceptualization, A.B.; project administration, A.B.; funding acquisition, G.S., L.C., A.B.

